# Representation of a Perceptual Bias in the Prefrontal Cortex

**DOI:** 10.1101/2023.07.27.550794

**Authors:** Luis Serrano-Fernández, Manuel Beirán, Ranulfo Romo, Néstor Parga

## Abstract

Perception is influenced by sensory stimulation, prior knowledge, and contextual cues, which collectively contribute to the emergence of perceptual biases. However, the precise neural mechanisms underlying these biases remain poorly understood. This study aims to address this gap by analyzing neural recordings from the prefrontal cortex (PFC) of monkeys performing a vibrotactile frequency discrimination task. Our findings provide empirical evidence supporting the hypothesis that perceptual biases can be reflected in the neural activity of the PFC. We found that the state-space trajectories of PFC neuronal activity encoded a warped representation of the first frequency presented during the task. Remarkably, this distorted representation of the frequency aligned with the predictions of its Bayesian estimator. The identification of these neural correlates expands our understanding of the neural basis of perceptual biases and highlights the involvement of the PFC in shaping perceptual experiences. Similar analyses could be employed in other delayed comparison tasks and in various brain regions to explore where and how neural activity reflects perceptual biases during different stages of the trial.

## Introduction

Perception is shaped by various factors including the current and previous stimulation, prior knowledge, and contextual cues [1], [2]. Collectively, these factors can contribute to the emergence of perceptual biases [2]–[10]. These biases manifest in various forms, such as the central tendency observed in tasks involving magnitude or duration reproduction [11]–[15], the contraction bias observed in delayed comparison tasks [7], [10], [16]–[21], and the phenomenon of serial dependence observed in delayed response experiments [22]–[24]. A normative Bayesian account posits that perception emerges through the integration of a noisy representation of the stimulus with prior knowledge about the sensory environment [1], [19], [25]. As the level of sensory noise increases, Bayesian observers tend to rely more heavily on their prior knowledge when forming their estimations. In certain situations, this can result in a perceptual bias towards perceiving stimuli parameters that are closer to the center of their distribution. For instance, this Bayesian framework has been suggested to explain the central tendency effect [26]–[29], the contraction bias [10], [19], [20], [30] and serial dependence [31]–[33].

Our current understanding of the relationship between neural activity and the contraction bias is still quite limited. However, there are some notable studies that have shed light on this topic [7], [20], [21]. One such study by Akrami et al. (2018) highlighted the importance of working memory in the manifestation of the contraction bias. Additionally, another study by Sarno et al. (2022) explored how the contraction bias modulates dopamine activity in the midbrain. Benozzo and Genovesio (2023) analyzed the activity of prefrontal neurons to investigate the effects of the contraction bias on the decision-making process. While these studies are intriguing, they do not address the issue of how Bayesian-like computations leading to the contraction bias manifest in the activity of cortical neurons.

Recently, some progress has been done in that direction. In tasks involving time interval reproduction, where subjects report their perception of duration, it has been observed that the curvature of state-space trajectories during the presentation of the sample interval is a distorted representation of the interval’s duration [14], [29]. This effect was also evident when the interval, instead of being reproduced, needs to be compared with another interval presented after a delay period [20]. Recurrent networks of spiking neurons trained on this task also displayed a curved manifold carrying a warped representation of the intervals during the presentation of the first stimulus. Throughout the delay period, the interval continued to be represented in a distorted manner by other geometric characteristics of the trajectories. Importantly, in both studies, these distorted representations aligned with the Bayesian estimation of the interval duration. This line of research provides valuable insights into the cognitive and neural processes involved in the formation of perceptual biases and their implications for decision-making. However, further studies are necessary to deepen our understanding of the specific computational mechanisms and population activity dynamics through which the biases emerge in cortical areas.

In this work, to investigate the neural representation of the contraction bias in cortical circuits and the underlying computational mechanisms, we examined the activity of prefrontal cortex (PFC) neurons during a vibrotactile frequency discrimination task performed by two monkeys [34], [35]. Our findings revealed that PFC neurons encoded information about the current stimuli but not about those in preceding trials. In contrast, we found evidence supporting the idea that PFC neural activity encodes aspects of the stimulus set’s structure, a basic requirement for Bayesian computations. To further investigate the compatibility of our findings with Bayesian principles, we developed a normative Bayesian model and successfully fitted it to behavioral data. Remarkably, the geometric characteristics of the state-space activity trajectories spanning from the presentation of the first frequency to the end of the delay period exhibited features that corresponded to the Bayesian estimator of that frequency. These results show that the dynamics of population activity in the PFC align with Bayesian concepts. This study opens the way for future research to investigate the underlying neural mechanisms and circuitry involved in the emergence of the contraction and other perceptual bias.

## Results

### Tasks

The research was conducted on recordings collected from two monkeys in previous experiments [34], [35] (Methods). Both monkeys had been trained to compare two frequencies, *f*_1_ and *f*_2_, separated by a delay period. However, while the first monkey was required to respond shortly after the offset of *f*_2_ (Fig. 1*A,* *Upper*), the second monkey had to wait for 3 s before communicating its decision at the moment of the Probe Up (PU) event (Fig. 1*A,* *Bottom*). Additionally, different stimulus sets were used; the first monkey was presented with the set 1, shown in Fig. 1*B,* *Left*, while the second monkey was presented with the set 2, depicted in Fig. 1*B,* *Right*. The probability distributions of *f*_1_ and *f*_2_ (Fig. 1*C,* *Left* and *Right*) are defined from the stimulus set. In both sets, frequency pairs (*f*_1_,*f*_2_) (stimulus classes) are organized along two diagonals, to be referred to as the upper and lower diagonals of the set. For later convenience classes were assigned a label (indicated inside the squares in Fig. 1*B*). Note that set 2 deliberately excluded any value of *f*_1_ that would allow subjects to predict the appropriate motor response.

**Fig. 1.**
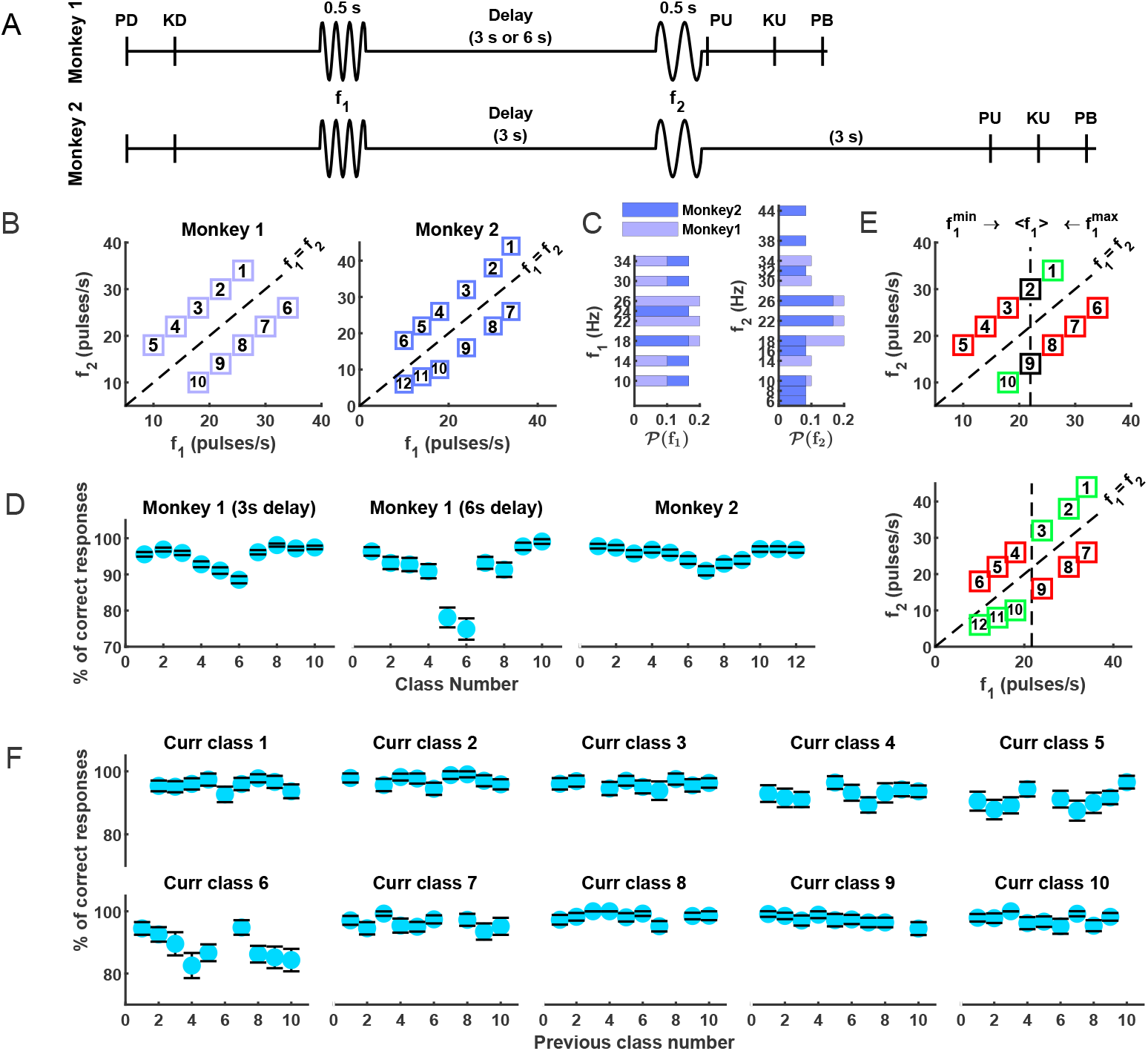
Task, stimulus sets and behavior. *(A)* Schemes with the timeline of the frequencies discrimination task for monkey one *(Upper)* and monkey two *(Bottom)*. A mechanical probe (Probe Down, PD) is applied to the smooth skin of a hairless finger on a restrained hand. To initiate each trial, the monkey presses a key with its free hand (Key Down, KD), indicating its readiness. A 500 ms long mechanical vibration (first stimulus, *f*_1_), followed by a delay period of 3 or 6 seconds, is then delivered. Subsequently, another 500 ms long mechanical vibration (second stimulus, *f*_2_) is applied. Once the mechanical probe is lifted (Probe Up, PU), the monkey is allowed to report its choice regarding the comparison between the two vibrations. At this point, the monkey’s free hand is no longer on the readiness key (Key Up, KU), and it proceeds to press one of two push-buttons (PB) to indicate its choice. The only difference between both schemes is a 3 s delay period between the offset of *f*_2_ and PU in the task of monkey two. *(B)* Stimulus set of possible classes or pairs (*f*_1_, *f*_2_) for monkey one (set 1; *(Left)*) and monkey two (set 2; *(Right)*). *(C)* 𝒫 (*f*_1_) and 𝒫 (*f*_2_) indicate the probability of the first *(Left)* and second *(Right)* frequency, respectively, as defined by the stimulus set for monkeys one (light purple) and two (dark blue). *(D)* Percentage of correct trials from monkey one with 3s delay *(Left)* and with 6s delay *(Middle)* and from monkey two *(Right). (E)* Contraction bias refers to the tendency of the first frequency (*f*_1_) to be coded in a contracted manner towards its distribution mean. This bias leads to errors in certain classes (red squares), while favoring others (green squares). *Upper* panel corresponds to set 1, while *bottom* panel corresponds to set 2. *(F)* Performance of monkey one is influenced by the previous trial. Each panel illustrates the accuracy of each current class as a function of classes in the previous trials. Data corresponding to the previous class being equal to the current class are omitted due to the small number of available trials; this low trial ratio stems from the recording method (see Methods). Error bars computed with 1,000 bootstrap resamples.

### Behavior. Monkeys exhibited the contraction bias

The percentage of correct responses (Fig. 1*D*) revealed that performance is contingent on the class, even though in either all (Fig. 1*D,* *Left* and *Middle*) or a majority (Fig. 1*D,* *Right*) of the classes the difference between the two frequencies presented in the trial possesses the same absolute value (Δ*f* = |*f*_1_ − *f*_2_| = 8 Hz). This discrepancy can be attributed to the presence of a contraction bias, which impacted decision accuracy based on the specific pairing of stimuli. This internal process caused a shift in the perceived frequency of the base stimulus *f*_1_ towards the center of the frequency range (⟨*f*_1_⟩ = 22 Hz and ⟨*f*_1_⟩ = 21.67 Hz, which are the means of *f*_1_ obtained from its probability distribution in Figs. 1*C,* *Left* and Fig. 1*C,* *Right*, respectively). Consequently, when *f*_1_ had a lower value, it was perceived as larger, whereas a higher value was perceived as smaller, as illustrated in Fig. 1*E*. Thus, the influence of the bias progressively hindered accurate evaluations for classes where “*f*_1_ *< f*_2_” as they were encountered from right to left (upper diagonals in Fig. 1*B*). In contrast, the bias increasingly facilitated classes where “*f*_1_ *> f*_2_” (lower diagonals). To uphold the bias structure, we designated the classes benefiting from the bias effects as positioned towards the extreme ends, while the central classes encountered restrictions imposed by the bias (Fig. 1*B*). In this classification arrangement, the classes positioned near the two ends, where the bias had a positive influence, demonstrated the highest level of decision accuracy. As already noticed in [10], when plotted against the class number, the resulting curves displayed V-shaped patterns, as depicted in Fig. 1*D*. We have also noticed that the performance in the current trial is influenced by the class presented in the previous trial, as it was observed in previous works [7], [36]. Specifically, when plotting the percentage of correct responses for the current trial class against classes in the previous trial, we noted that the performance is not constant (Fig. 1*F*).

In this work we will focus on the effect that prior knowledge combined with observation on the current trial may have on the occurrence of the contraction bias. The Bayesian explanation predicts that increasing the length of the delay period will lead to an increment in the magnitude of the bias [23]. This is because the maintenance in short-term memory of *f*_1_ may lead to greater uncertainty about its value. To verify this prediction, we started by quantifying the contraction bias *B* as the root-mean-square deviation of the animal performance (Methods). Calculating *B* for two different durations of this interval, Δ = 3 s or Δ = 6 s (monkey one), yielded *B* = 0.036 ± 0.0035 and *B* = 0.078 ± 0.0086, respectively, confirming the Bayesian prediction. A Mann-Wilcoxon test assessed that both distributions (obtained from a bootstrap procedure) were significantly different (p-value=0).

### PFC neurons carried information about the current but not about the previous trial *f*_1_

A possible source for the contraction bias is sensory history [7], [20], [36], acting, e.g., through the persistence in the population firing activity of the representations of stimuli applied in previous trials. However, this does not imply that the activity of neurons in a given area should show such dependence, even if the neural activity in that area exhibits the contraction bias in some manner. This is because of the potential involvement of multiple brain regions in generating the contraction bias [7], [30]. In the posterior parietal cortex (PPC) of rats, neuronal activity during the delay period lacks mutual information (MI) about the current stimulus. However, during the inter-trial interval, it did exhibit MI concerning the previous trial’s *f*_1_ [7]. Meanwhile, PFC neurons, crucial for working memory, might contribute to the contraction bias through Bayesian computations akin to those proposed in [19]. Moreover, PFC neurons could potentially receive information from PPC about short-term stimulation history. This suggests that these two areas could potentially form a cortical circuit generating the contraction bias and associated properties [37]. However, this presumption is not definitive due to the distributed nature of working memory [38]; other brain regions might also participate in this circuit. Therefore, the transmission of previous trials sensory information from PPC to PFC neurons might not be mandatory.

To clarify the behavior of PFC neurons, we first calculated the MI that each neuron carried about the current trial *f*_1_ value and its value from the previous trial (Methods), using correct and fixed choice current trials (to remove the confounding factors of reward and action). Note that then the MI about the current trial *f*_1_ is equivalent to the MI about the current class. Subsequently, we determined the fraction of neurons for which those pieces of information were found to be significant. In both monkeys, for the two choices, neurons exhibited significant information regarding the class in the current trial (SI Appendix, Fig. S1). The population size conveying this information initially increased upon presentation of *f*_1_, then decreased during the early phase of the delay period, and eventually recovered prior to the onset of *f*_2_ (Fig. 2*A,B*, blue lines). However, there was no population of neurons carrying significant information about the value of *f*_1_ from the previous trial (Fig. 2*A,B*, red lines). This result is particularly evident for monkey two, as the proportion of neurons exhibiting MI about the current trial becomes nonsignificant already during the second delay period of the task (Fig. 2*B*). If analogous analyses are conducted for the information concerning *f*_2_, one would obtain very similar results (SI Appendix, Figs. S2 and S3). This is because, by fixing the outcome and the choice in a trial, knowledge of *f*_1_ inherently implies knowledge of *f*_2_.

**Fig. 2.**
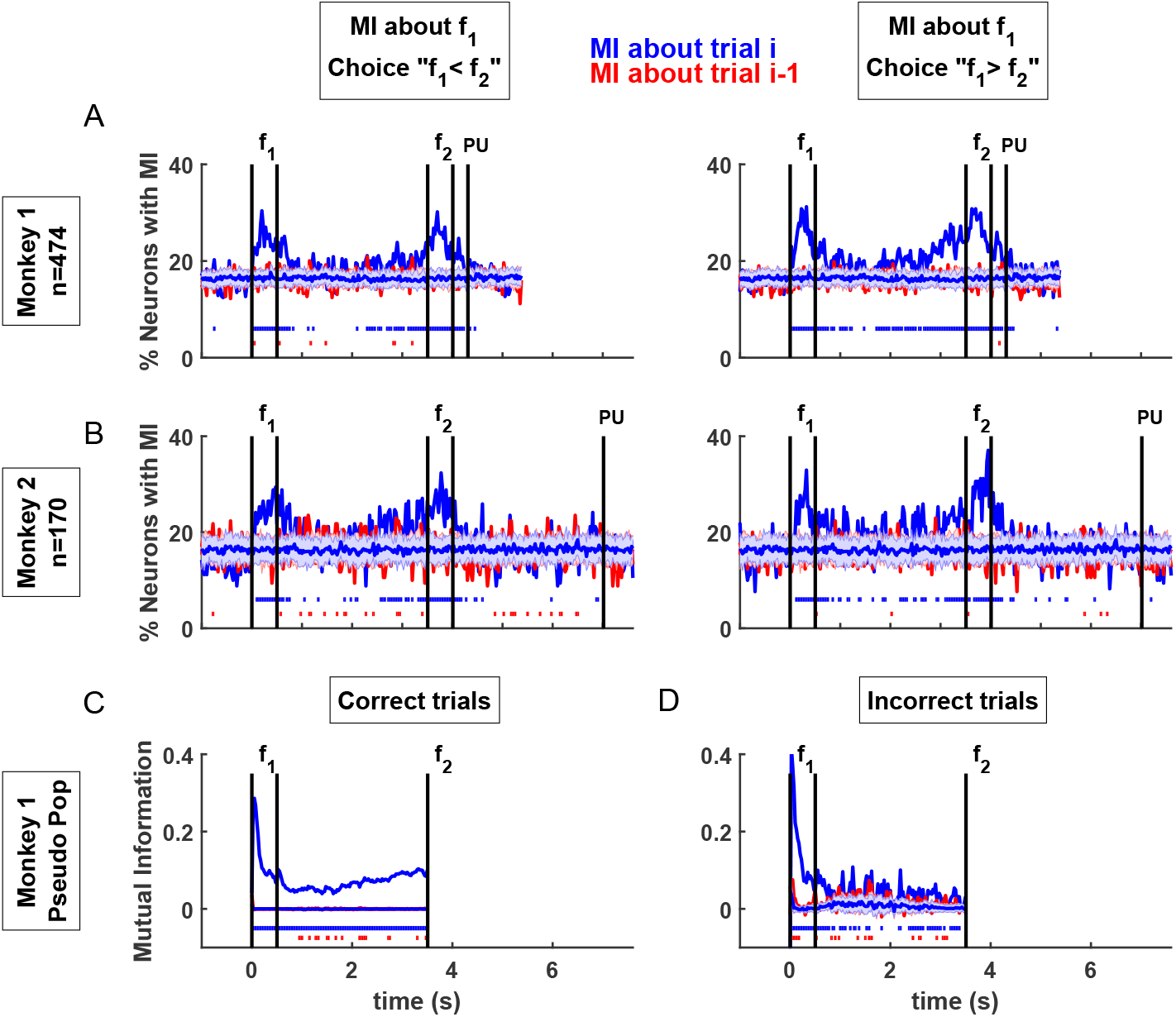
PFC neurons exhibit significant mutual information (MI) about current trial *f*_1_ but not on the previous trial *f*_1_. *(A)* Percentage of neurons from monkey one with significant MI about the current (blue line) and previous (red line) first stimulus, conditioning on correct an fixed choice (*Left*: choice “*f*_1_ *< f*_2_”; *Right*: choice “*f*_1_ *> f*_2_”) current trials. Note that this effectively fixes the class in the current trial. Horizontal blue and red shadings refer to the mean and standard deviation computed from shuffled data (see Methods) for current and previous trial, respectively. The horizontal lines of dots at the bottom indicate runs of consecutive time bins where a significant percentage of neurons exhibit significant MI, assessed through run length analysis (see Methods). *(B)* Same MI analysis for monkey two. *(C-D)* MI analysis using the z-score distribution with correct *(C)* and incorrect *(D)* trials from all neurons tuned to *f*_1_ in the dataset of monkey one. In this analysis trials with both choices were included for each *f*_1_.

Calculating MI for stimuli in error trials poses a challenge due to their low occurrence, even in classes negatively affected by the contraction bias (Fig. 1*D*). To determine if there is information regarding *f*_1_, whether in the current or preceding trial, we employed a methodology detailed in [39]. This approach hinges on the conditional distributions of z-scores from all neurons in the database tuned to *f*_1_ (Methods). Additionally, in this analysis, we did not differentiate trials based on choice. In error trials, our analysis revealed that this population of neurons conveyed MI about the value of *f*_1_ in the current trial (Fig. 2*D,* *blue line*), aligning with the findings in the medial premotor cortex as reported by Vergara et al. (2016) [39]. However, we did not observe any MI concerning the value of *f*_1_ presented in the preceding trial (Fig. 2*D,* *red line*). When applying the same methodology to correct trials, our results are consistent with those presented in Fig. 2*A*: there exists MI about the value of *f*_1_ in the current trial (Fig. 2*C,* *blue line*), while there is no information about its value in the previous trial (Fig. 2*C,* *red line*).

These analyses specifically pertain to the transient sensory history and do not rule out the possibility of PFC neurons possessing information about the stationary stimulation history.

### PFC neurons reflect aspects of the set structure

Earlier we saw that, in agreement with the Bayesian hypothesis, the contraction bias *B* increases with the duration of the delay period. We now turn our attention to examining the extent to which the activities of neurons in the PFC reflect features about the structure of the stimulus set, before *f*_2_ is presented. We start by acknowledging that within set 1 (Fig. 1*B,* *Left*), classes 4-7 can be uniquely determined by the value of the first frequency. We then investigated the presence of neurons whose activity, before the presentation of *f*_2_, is correlated with the choice made at the end of the trial. To assess the presence of these neurons in the data collected from monkey one, we evaluated the choice probability index (CPI) [40], [41], a metric derived from the firing activity distributions in correct and error trials within a specific class, providing an estimate of the probability of decoding the choice from neural activity (Methods). A CPI of 0.5 signifies complete overlap of the two distributions. A CPI of 1 or 0 indicates entirely distinct distributions, whereas a CPI less than 0.5 indicates that the difference between the two distributions is opposite to their expected outcome. Because of the restricted number of error trials, we computed the CPIs using the set of z-scores from all neurons in the dataset tuned to *f*_1_ (Methods).

For all the classes determined by their value of *f*_1_ (classes 4-7), the CPIs turned out to be significant before the presentation of the second frequency. In classes 6 and 7 (where the correct choice is “*f*_1_ *> f*_2_” and rewarded trials tend to have firing rates larger than incorrect ones), the significant CPI values were greater than 0.5, while for classes 4 and 5 (where the correct choice is “*f*_1_ *< f*_2_” and rewarded trials tend to have firing rates less than incorrect ones), they were less than 0.5 (Fig. 3). This supports our hypothesis that in classes 4-7, PFC neural activity encodes the choice prior to the presentation of *f*_2_, indicating that it has captured elements of the stimulus set’s structure. Since the CPI is significant not only for classes 5 and 6 but also for classes 4 and 7, this finding eliminates the possibility of attributing the ability to predict the choice to *f*_1_ values being on the boundary of its range. In the remaining classes (1-3 and 8-10), we expect non-significant CPIs as it is not feasible to predict the choice in these classes before the presentation of *f*_2_. Regrettably, the absence of enough error trials in those classes prevented us from confirming this assertion. The above arguments assume that the CPI of PFC neurons is primarily influenced by top-down processes [42], with minimal or no contributions from stimulus fluctuations (*f*_1_). This view is supported by the observation that the CPI is rarely significant during stimulus presentation. In contrast, in a classic experiment, medial temporal neurons exhibited a weak but significant CPI during stimulus presentation, while visual evidence is being accumulated [40]. A modeling study considering both bottom-up and top-down effects on choice indicated that the contribution of stimulus fluctuations decreased during the stimulation period, whereas the top-down contribution remained sustained after the stimulation period [43]. If applicable to our situation, this would imply that the significant CPI observed during the delay period (Fig. 3) is not due to fluctuations in the first frequency, suggesting that PFC neuron activity reflects an aspect of the stimulus set.

**Fig. 3.**
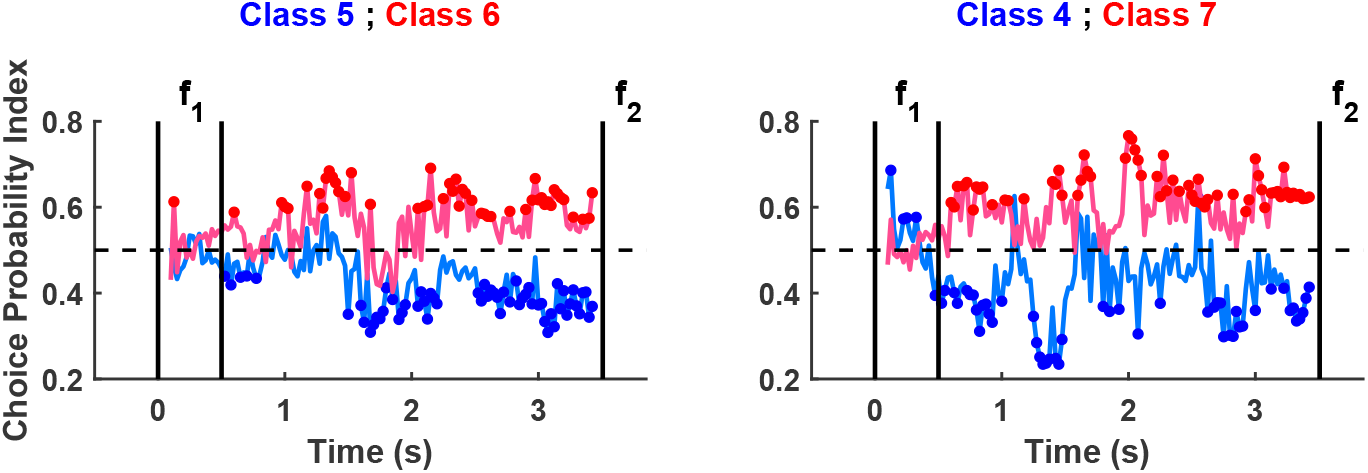
PFC neural activity reflects the structure of the stimulus set. Temporal evolution of the Choice Probability Index (CPI) from the onset of *f*_1_ until the onset of *f*_2_ using data collected from monkey one. The analyzed classes were those determined solely by their *f*_1_ value: classes 5, 6 (*left*), 4 and 7 (*right*). Solid lines represent CPI dynamics over time using z-score distributions from all neurons tuned to *f*_1_. Colored dots denote significant CPI values (see Methods).

### State-space Trajectories of Neural PFC activity

Our approach is based on the understanding that neural circuits encode and process information by utilizing the dynamics of neural populations within a state-space framework [5], [20], [44]–[49]. Then, to explore the relationship between behavior and neural firing activity, we conducted our analysis at the population level, using all the recorded neurons, and projected the population activity (z-score) onto the state-space directions that accounted for a minimum of 90% of the data variance (Methods).

Our findings revealed that in correct trials the population z-score evolved separately from each other during both the presentation of the *f*_1_ (Figs. 4*A,D*) and the delay period (Figs. 4*B,E*). About one second after the offset of the first stimulus, the trajectories displayed reduced velocity. However, by the end of the delay period, there was a slight increase in velocity, better appreciated in Figs. 4*B*, and 4*E*. The trajectories reached terminal states dependent on the value of *f*_1_, with these states notably exhibiting an organized alignment (see the magenta line in Figs. 4*B,E*). As the delay period neared its end, there was an increase in the percentage of variance accounted for by the first principal component (PC) (Figs. 4*C,* *Right*, and 4*F, Right*), while the subsequent PC decreased. For these reasons we considered the set of terminal states as a relevant geometric characteristic of the trajectories (magenta lines in Figs. 4*B,E*). The striking similarity in the trajectories observed in both monkeys (as depicted in Figs. 4*A,D*, and Figs. 4*B,E*) strongly supports the notion that the corresponding PFC neural populations are engaged in similar computational processes.

**Fig. 4.**
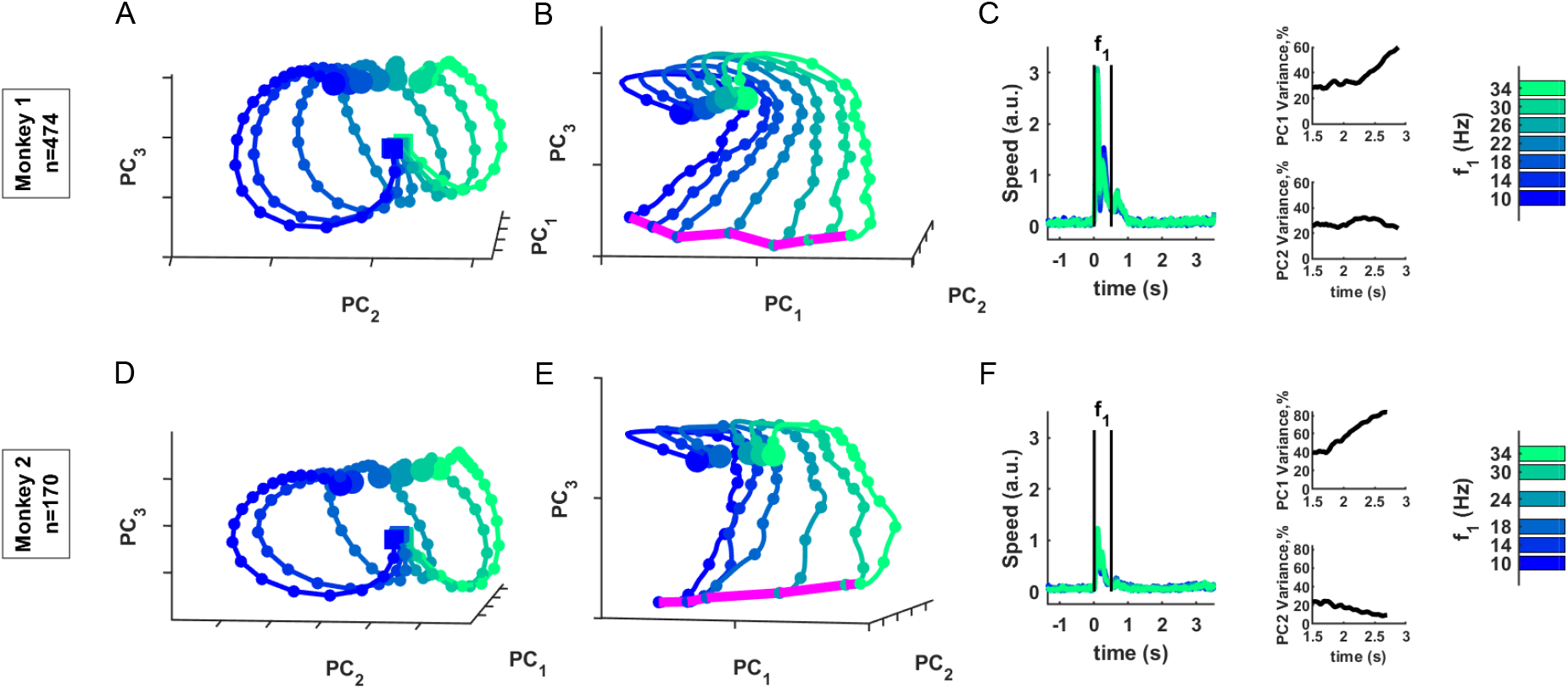
State-space trajectories and terminal states. *(A-C)* Monkey one. *(A)* Projection of neural trajectories onto the first three PCs during the presentation of *f*_1_. Squares and big circles represent the onset and the offset of *f*_1_, respectively. Small circles indicate neural states every 25 ms intervals. It is important to note that the distance between consecutive small circles reflects the speed of neural trajectories. *(B)* State-space neural trajectories during the 3 s delay period. Big circles indicate the offset of *f*_1_. Small circles represent neural states at 250 ms increments. Magenta line connects the terminal states at the end of the delay period. *(C, Left)* Speed of neural trajectories over time from 1.5 seconds before *f*_1_ until the end of the delay period. *(C, Right)* Percentage of variance explained by the first (*Upper*) and second (*Bottom*) PCs at the end of the delay period; both variances sum around 90%. *(D-F)* Same state-space analysis but for monkey two. Only correct trials were considered.

Nonetheless, the current finding does not specify which estimator of *f*_1_ is upheld by the network during the period starting from the presentation of *f*_1_ until the end of the delay interval. To investigate this matter and determine the geometric properties of the trajectories that encode the estimator, it is essential to establish a link between behavior and population activity. This aspect will be the focus of our upcoming analysis.

### Bayesian Model and Fits of Behavioral Data

By leveraging the benefits of normative models in identifying underlying network computations associated with observed behavior, we developed a Bayesian model that incorporates detailed *a priori* knowledge (Methods. Prior Bayesian proposals and models are detailed in previous works [19], [30]). We adopted the perspective of an observer who makes imperfect measurements of the frequencies *f*_1_ and *f*_2_, possesses knowledge of the stimulus set (including the mean value and variance of the *f*_1_ distribution), and can formulate beliefs regarding the presented stimuli. We assumed a Gaussian nature of the frequency measurements, with means *f*_1_ and *f*_2_, and standard deviations *σ*_1_ = *W*_1_*f*_1_ and *σ*_2_ = *W*_2_*f*_2_, respectively. Here, *W*_1_ and *W*_2_ represent the corresponding Weber fractions.

In our model, in accordance with the results in Fig. 3, we assumed that the observer possesses good knowledge of the stimulus set. Note that, despite having demonstrated the absence of transient sensory history effects in PFC neural activity (Fig. 2, red lines), this does not rule out the possibility that neurons in this area may compute (or receive from elsewhere) the long-term sensory history of the first frequency. Indeed, previous models of the contraction bias have incorporated a dependence on the mean of the stimulus set [7], [17], [18], and experimental findings indicate that this dependence operates within a Bayesian framework [30]. We then assumed that the decision in the current trial *n* was based on an observation comprising a mixture of the noisy measurement of the first frequency 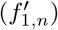, along with an estimate of the stationary component of the sensory history of this frequency. From a mathematical standpoint, the observation (*o*_1,*n*_) consists in the combination of 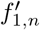 with an online estimate of the long-term (stationary) sensory history (*o*_1,*n*−1_),

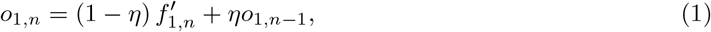

where if *η* = 0 there is no contribution from the long-term sensory history while if *η* = 1 the current stimulus does not contribute. In the Gaussian approximation, the likelihood *P* (𝒪_1_|*f*_1,*n*_) (𝒪_1_ is the stochastic Gaussian variable associated to the observation in the current trial, and *f*_1,*n*_ is the first frequency presented in that trial) has the mean and variance (see Methods for details)

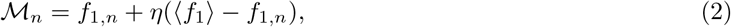

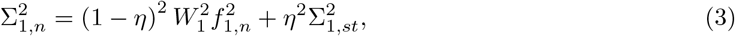

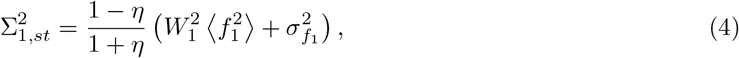

where 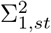 is the variance of the stationary distribution of observations of the first frequency (see SI Appendix for mathematical derivation) and 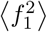 and 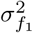 are the second moment and variance of in the first frequency in the stimulus set. It should be noted that both ℳ and Σ_1_ play a role in the contraction bias phenomenon. The mean explicitly expresses how *f*_1,*n*_ is pulled towards ⟨*f*_1_⟩, and this contraction is proportional to the difference between the actual value of the frequency and its mean. Qualitatively, a Bayesian contribution to the contraction bias occurs when the prior has a more substantial impact on the posterior of the first frequency compared to the second frequency [19], particularly when Σ_1_ exceeds *σ*_2_. The priors are defined as the probabilities of the stimuli (Fig. 1*C*) and correspond to the inherent structure of the stimulus set (Methods). Finally, we assumed that *f*_2_ is solely affected by measurement noise.

To fit the model parameters (*η, W*_1_ and *W*_2_), for each monkey we considered the corresponding accuracy curve (Fig. 1*D*), and used it to define the similarity between model and data (Methods). The fits and the optimal parameter values for the two animals are shown in Fig. 5. The data recorded in monkey one from two different delay period durations allowed us to assess the significance of *η*, the parameter linked to long-term sensory history. To do this, we compared two fits of the accuracy data for the 6-second delay period. In one fit, all three parameters were optimized (Fig. 5, *Middle*), while in the other, only *η* was allowed to vary, with the other two kept at the values obtained for the 3-second delay period data (Fig. 5, *Left*). If the sole purpose of *η* were to accommodate the long-term sensory history, one might anticipate that the values of *η* obtained in the two fits would be similar. The comparison of the two fits indeed confirmed this expectation, with *η* changing by less than 1% (SI Appendix, Table S1). Given the expected influence of the delay period duration on the noise parameter *W*_1_, we extended our analysis to optimize both *η* and *W*_1_, while keeping *W*_2_ fixed as before. Once more, *η* exhibited minimal alterations, while the Root Mean Square Error (RMSE) of the fit differed by only 3.5 % compared to the initial fit (SI Appendix, Table S1). Finally we compared the model fits with those from a model where *η* = 0 [19], noting that AIC and BIC tests favored the former fits (SI Appendix, Table S2).

**Fig. 5.**
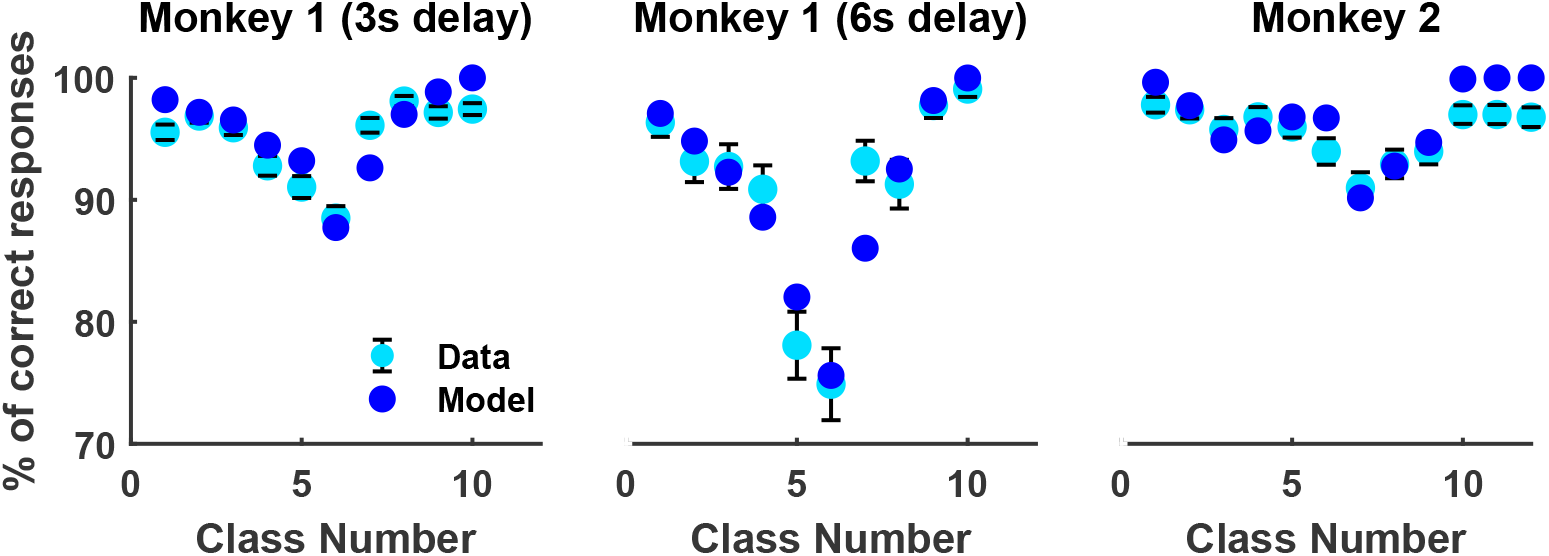
Bayesian fits of the Accuracy curves. Cyan dots with error bars correspond to behavioral data while dark blue circles represent the fit using Eqs. (14-15). *(Left)* Monkey one with a 3-second delay interval. The adjusted model parameters are the Weber fractions of the first (*W*_1_ = 0.064) and second (*W*_2_ = 0.088) frequency and the sensory history parameter (*η* = 0.509). *(Middle)* Monkey one with a 6-second delay period. The fitted parameter values are *W*_1_ = 0.057, *W*_2_ = 0.110 and *η* = 0.634. *(Right)* Monkey two. The fit parameters are *W*_1_ = 0.054, *W*_2_ = 0.108 and *η* = 0.430. Note that the relevant quantity to determine the Bayesian contribution to the bias is not *W*_1_ but Σ_1_. In both monkeys, the condition Σ_1_ *> σ*_2_ was satisfied for the majority of the frequency pairs (*f*_1_, *f*_2_).

It’s important to note that accuracy behaves differently along the two diagonals, with its minimum specifically occurring in the first class of the second diagonal (Fig. 5. Class 6 in set 1 and class 7 in set 2). Our model provides an explanation for this phenomenon. Essentially, Weber’s law introduces an asymmetry in the bias that can change the location of the minimum: in set 1 the standard deviation Σ_1_ in class 6 is larger than in class 5 (Eq. 3), leading to a greater bias in the first of these classes and, consequently, poorer performance.

In the model we assumed that the Bayesian observer has a good understanding of the experimental set, particularly the transition probabilities. To test the validity of this assumption, we extended the model by introducing non-zero probabilities (*ϵ*) for transitions not allowed by the stimulus set and we fitted the parameters of the extended model (Methods). Confirming our assumption, *ϵ* turned out to be quite small for monkey 1 and reasonably small for monkey 2 (SI Appendix, Table S3) while the values of the other parameters hardly change.

### Relationship Between PFC State-Space Geometry and Behavior

We are now investigating the link between state-space geometry and behavior. In this task, the neural activity is expected to represent an estimate of the frequency *f*_1_ during its stimulation and maintain a representation throughout the delay period until its conclusion. Consequently, we anticipate that during these epochs, an estimate of *f*_1_ may be reflected in the geometric properties of the trajectories [14], [20], [29]. Our observations indicate that during the presentation of the stimulus, the trajectories evolve separately while maintaining finite relative distances (Fig. 4*A,D*), during the delay period still remain distant from each other (Fig. 4*B,E*), ultimately converging to a line of terminal states at the end of the delay epoch (Fig. 4*C,F*). Based on these findings, we hypothesize that estimations of *f*_1_ are represented: firstly, in the relative distances between trajectories as they evolve under the influence of the first stimulus, then, during the delay period, in the relative distances maintained by the trajectories, and finally, in the distances between consecutive terminal states.

However, the specific nature of the estimator used by the cortical network remains unknown, as only the comparison choices are reported by the animals in the discrimination task. This differs from magnitude reproduction tasks, where the reproduced magnitude can be considered as a noisy version of the reported estimate [28]. Due to the limited information regarding the employed estimator for *f*_1_, we explored three alternatives: the true *f*_1_, the Bayesian estimator (*f*_1,*Bayes*_), and the Maximum a Posteriori (MAP) estimator (*f*_1,*MAP*_) (Methods). We examined how these estimators were associated with the relative distances between neural trajectories during the stimulation and short-term memory periods, as well as with the distances between terminal states, at the end of the delay epoch.

To evaluate the estimator that most accurately captured the relative distances between neural trajectories during the stimulus presentation phase, we employed the following procedure. We focused first on monkey one during the presentation of the first stimulus. We computed the relative distance 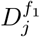 (t) between each trajectory (indexed as *j* = 1, …, 7) and the reference trajectory (*f*_1_=22 Hz) using the KiNeT methodology [13] (Methods). The temporal evolution of the distances between trajectories (Fig. 6*A*) shows that the distance increases less when the frequency is further away from the reference trajectory. As a consequence, when plotting the average distance between trajectories during the stimulus presentation as a function of the stimulus value (Fig. 6*A* inset), we observe an encoded sigmoid-like warping of the stimulus. We subsequently investigated whether the distance 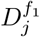 more effectively encoded the true *f*_1_ values, the Bayesian estimator of *f*_1_, or the MAP estimator. To compare these alternatives, we individually fitted a linear expression 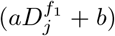 to each of these three estimates (Fig. 6*B*) and assessed their RMSEs [14] (Fig. 6*C*). The results strongly supported the hypothesis favoring that the distances between trajectories represent a Bayesian estimator of the stimulus. Subsequently, we conducted a comparable analysis focusing on the relative distances during the delay period, denoted as 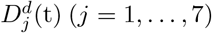 We found again that the relative distances are warped or contracted towards the reference trajectory (Fig. 6*D*). Analogously, the obtained results provided support for the hypothesis that such geometry of neural trajectories favors the Bayesian estimator (Fig. 6*D-F*). Finally, we applied a similar analysis considering the set of states of neural trajectories at the end of the delay period which lie within a low-dimensional space (where two dimensions explain 84% of the variance). Then, we conducted a comparative assessment of the distances *δ*_*j*_ (between each final state and the one with the smallest *f*_1_) with the true *f*_1_ value and the Bayesian and MAP estimators. Once again, the results strongly supported the Bayesian estimator (Figs. 6*G-I*). Collectively, the outcomes of these three tests consistently suggest that the representation of the first frequency is effectively captured by its Bayesian estimator. This conclusion was further supported by conducting a similar analysis on the data from monkey two (SI Appendix, Fig. S4).

**Fig. 6.**
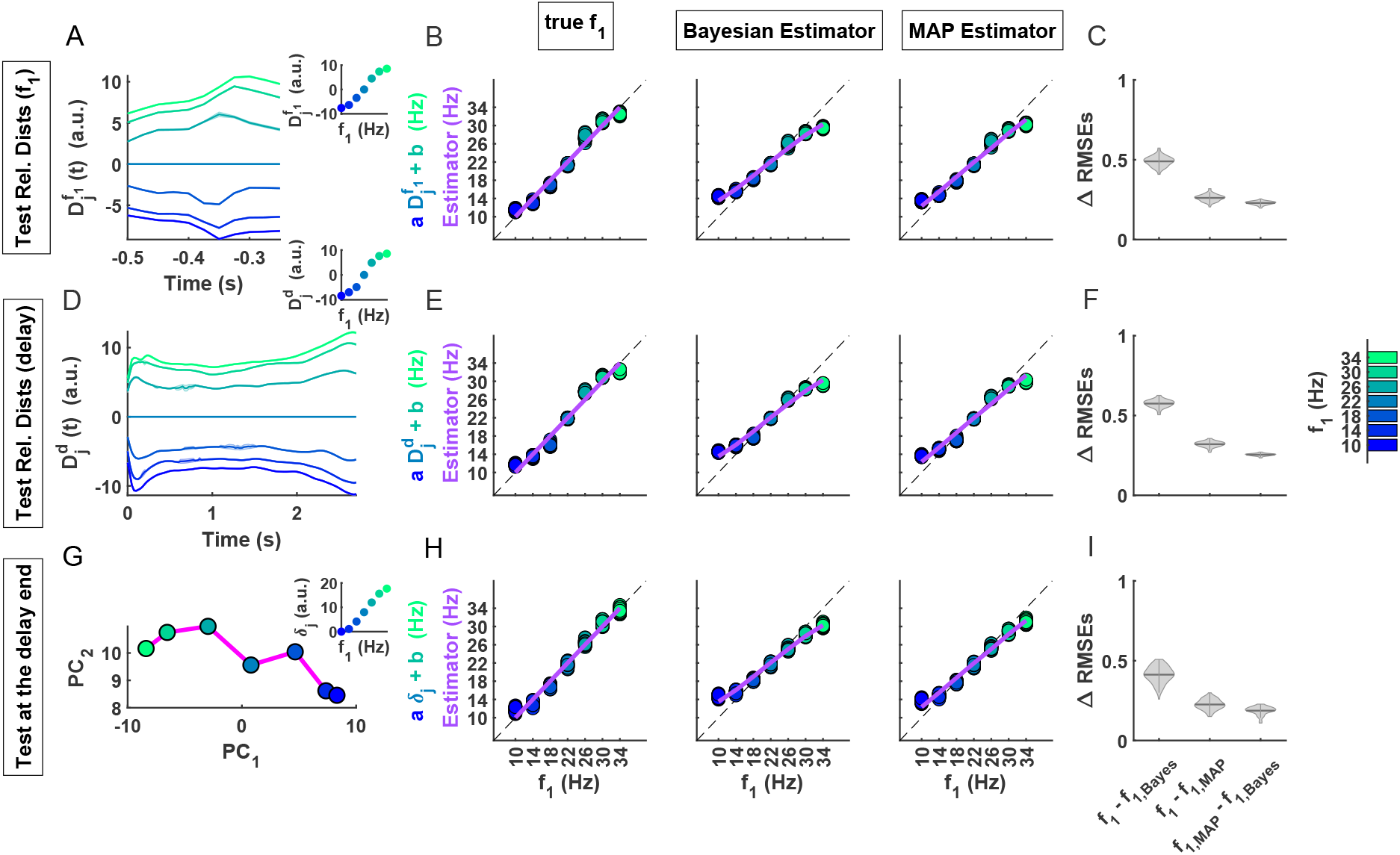
Several geometric features from monkey one code the Bayesian estimator. *(A-C)* KiNeT trajectories and activity-behavior relationship during the presentation of the first frequency. *(A)* Relative distance, 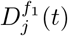, between each trajectory *j* = 1, …, 7 and the reference trajectory *f*_1_=22Hz using KiNeT analysis. Alignment is located at the offset of the first stimulus. The state space that embeds these neural trajectories is constructed using 6 PCs, which account for 92% of the variance. Shadings were built from the mean ± 95% CIs of 100 bootstrap resamples. *Inset* displays *D*_*j*_, the temporal means of relative distances as function of *f*_1_. *(B)* Linear regression of the mean relative distances 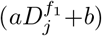 to the true *f*_1_-value *(Left)*, the Bayesian estimator *(Middle)* and the MAP estimator *(Right)* as a function of *f*_1_. Colored dots represent multiple estimates of the regression obtained through bootstrapping (100 resamples). Purple solid lines indicate the true *f*_1_ (*Left*), its Bayesian estimator *(Middle)* and its MAP estimator *(Right)*. The unity line is represented by a black dotted line. *(C)* Violin plots display the empirical probability density of ΔRMSEs on the y-axis, which represents the differences in RMSEs between the three hypotheses (*f*_1_, Bayesian estimator, and MAP estimator) indicated on the x-axis. A positive ΔRMSE value indicates that the second of the two estimators fits the geometric feature better. The Bayesian estimator fits the data better than the alternative hypotheses for all tests. The width of the gray area represents the density of the shufflings in that range. The horizontal line within each violin plot indicates the mean of the distribution. *(D-F)* KiNeT trajectories and activity-behavior relationship during the delay period. Same format as *A-C*, as well as same alignment at the offset of *f*_1_. Activity analysis with KiNeT was build from a state space of 30 PCs explaining 90% of variance. Again, Bayesian hypothesis is chosen by the 100 resamples. *(G-I)* Terminal states and activity-behavior relationship at the end of the delay period. *(G)* Line of terminal states (magenta line and circles) embedded within a 2-dimensional subspace (84% of variance). *Inset* shows the Euclidean distances along the magenta line, *δ*_*j*_, between each *f*_1_ terminal state and the smallest *f*_1_ (taken as reference), as function of *f*_1_. *(H-I)* activity-behavior relationship in the same format as *B-C*. One more time, Bayesian estimator is preferred.

It is noteworthy that the dynamics of population activity directly transformed the true value of *f*_1_ into its Bayesian estimator. The computations necessary to integrate prior knowledge and current sensory information were not explicitly carried out. This phenomenon resembles what is observed in time interval estimation experiments [14], [29] and in simulated recurrent neural networks trained to discriminate between the durations of two time intervals [20].

## Discussion

Previous studies have suggested that the contraction bias, a phenomenon in perception, can be attributed to the integration of prior knowledge with noisy sensory observations [10], [19], [20], [30], [50]. In the current work, we aimed to investigate the validity of this proposition and explore how Bayesian integration manifests in the population activity of PFC neurons. To accomplish this, we examined neural recordings from monkeys engaged in a discrimination task involving two tactile frequencies within the flutter range [34], [35]. Throughout the task, we observed distinct geometric characteristics in the state-space trajectories, which consistently contracted the representation of the first frequency, depicting low frequencies with values higher than their actual values and high frequencies with values lower than their true counterparts. This contraction was evident during both the stimulus presentation and the subsequent delay period, as reflected in the relative distances between trajectories. Towards the end of the delay period, these contractions converged towards a line of terminal states. Importantly, these contractions conform with the predictions of a Bayesian model, indicating that they underlie the manifestation of the contraction bias in the activity of PFC neurons.

Our findings can be compared to recent experimental and computational studies investigating neural activity in other tasks that exhibit similar perceptual biases. In a temporal reproduction task, it was found that a curved one-dimensional manifold exhibited a warped representation of the time interval, which was consistent with the distortion predicted by its Bayesian estimator [13], [14], [29]. Computational studies have demonstrated that the contraction bias can also occur in time interval discrimination tasks [20]. To investigate the underlying mechanisms of this bias, the authors trained recurrent neural networks composed of spiking neurons on such tasks. In the trained networks, the bias arose from the integration of prior knowledge and the current observation. Interestingly, similar to the findings in the temporal reproduction task [29], a curved one-dimensional manifold displayed a contracted representation of the first time interval, aligning with the predictions of the Bayesian estimator. Moreover, during the delay interval, the state-space trajectories exhibited similar contraction in their relative distances. In alignment with another modeling study [37], we encountered difficulties in effectively fitting the behavioral data with the Bayesian model in [19]. However, our Bayesian model, integrating the stationary sensory history, showed a better fit and received support from the model comparison tests. Although the current analysis in the PFC does not include a study on how artificial spiking recurrent neural networks might solve the tactile frequency discrimination task, there are now techniques available to train these networks either by abstractly implementing the task [51] or by directly using experimental data [52].

An intriguing finding emerged from our study, revealing that neurons in the PFC encode information specifically about the current first stimulus, while remaining insensitive to stimuli presented in previous trials, as observed across both correct and incorrect trials. This stands in stark contrast to observations in the posterior parietal cortex (PPC) of rodents, where the opposite pattern was observed [7]. The absence of short-term sensory history information in the PFC despite its presence in the PPC might be attributed to the highly distributed nature of working memory [38]. Additionally, our data favored a parameter linked to the long-term sensory history. The contraction bias may manifest in various cortical areas in different ways, which could reflect how it is constructed and expressed in behavior. To gain a comprehensive understanding of the emergence of this bias, it is crucial to extend the current study to other implicated brain areas and examine the relationship between neuronal population activity and behavior in each case. Expanding our study in this way would provide a comprehensive understanding of this bias and shed light on the distributed nature of working memory. A limitation of our study is the small number of simultaneously recorded neurons, which limits our ability to test predictions of the Bayesian framework that require analyzing state space trajectories in single trials [29]. Specifically, we cannot assess whether the variance of the relative distances, either during the presentation of the first frequency or during the delay period, is lowest for values of *f*_1_ at the extreme of its distribution and largest at its mean. Conducting an experiment to directly test this prediction would be interesting.

Despite not finding a direct encoding of the previous trial in the recorded PFC activity, the prior is consistently carved into the geometry of neural trajectories during the stimulus presentation. A recent study in monkeys performing a discrimination task in contrast has shown that a bias towards the choice and outcome of only the previous trial is reflected on the neural activity of the current trial, while other behavioral biases have a timescale of tens to hundreds of trials [53]. It remains unclear how different perceptual and behavioral biases are integrated at different timescales. Future work, with carefully designed trial correlations and recordings over training, shall clarify the temporal processing of different biases at the neural and behavioral level.

In contrast to a previous study focusing on the decisional stage of PFC neurons [21], our research investigates the earlier stages of the task, namely the stimulation and maintenance epochs. Our findings reveal that the Bayesian estimator is consistently reflected in the state-space geometry from the presentation of the first stimulus until the end of the delay period. During these task epochs we persistently noticed the influence of prior knowledge shaping a low-dimensional manifold. The associated dynamics directly translated *f*_1_ into its Bayesian estimator. Interestingly, this pattern was evident in both monkeys, even though they were trained with different stimulus sets. Our findings, alongside previous studies involving time reproduction tasks in monkeys [29], propose a broader computational framework for Bayesian integration. This framework implies that diverse geometric attributes, like curvature in timing tasks and relative distances in discrimination tasks, contribute to crafting a contracted representation of the stimulus and shape perceptual biases. This challenges the hypothesis that Bayesian computations are obtained by sequentially combining prior information with the current stimulus but aligns with the findings observed in other tasks [54]. Furthermore, our investigation of PFC neural activity demonstrates that the contraction bias is already evident upon the presentation of the first stimulus, suggesting that it may originate from a sensory area where it begins to take shape. This observation is in line with the findings of Preuschhof et al. (2010) [30], supporting the integration of prior information in early brain areas such as S2. Overall, our study provides new insights into the temporal dynamics and neural mechanisms underlying the contraction bias, contributing to a deeper understanding of this phenomenon.

## Methods

### Experimental Data

Data from two monkeys, previously trained to perform a somatosensory vibrotactile frequency discrimination task [34], [35], [55], were utilized in this study. The monkeys, named RR13 and RR15 (monkey one and monkey two, respectively), underwent training on distinct sets of vibrotactile frequency classes. The task was executed with a high level of accuracy, and our analysis specifically focused on correct trials. Both monkeys were recorded with a 3-second delay interval, while additional recordings were obtained from monkey RR13 with a 6-second delay interval. However, only monkey two had to wait for 3 s before reporting its decision. 474 neurons from monkey one and 170 cells from monkey two were considered for this study. Monkey one neurons were recorded over 116 experimental sessions while monkey two units proceeded from 56 sessions. In each session, there were around 10 blocks of trials. Each block had as many trials as there are classes in the set and classes were randomly chosen without replacement. A total of 10,540 trials of neural recording were considered for monkey one and 5,972 trials for monkey two.

### Data analyses: Behavior

#### Accuracy curve

In order to examine the discrimination properties, we plotted the percentage of correct trials as a function of the index of enumerated classes (Fig. 1*D*). *Phenomenological measure of the contraction bias*. Denoting the probability of deciding “*f*_1_ smaller” (*S* ≡ “*f*_1_ *< f*_2_”) given a presented class *C* as *P* (*S*|*C*), a phenomenological measure *B* of the contraction bias was defined as the root-mean-square deviation of *P* (*S*|*C*) on all classes, relative to classes 3 and 8 in case of monkey one (Fig. 1*B,* *Left*)

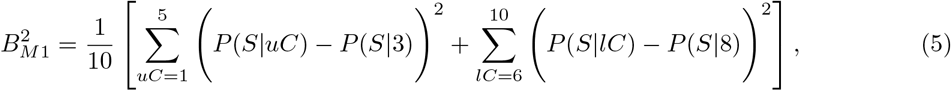

and relative to classes 3 and 9 in case of monkey two (Fig. 1*B,* *Right*)

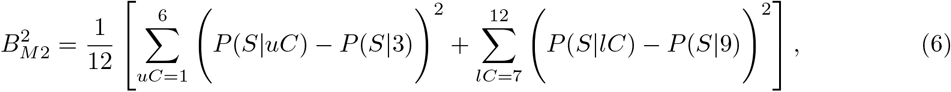

where *uC* and *lC* denote classes on the upper and the lower diagonals of the stimulus set, respectively. Note that the value of *B*^2^ corresponding to the minimum possible bias equals 0, while for the maximum possible bias it is 3*/*5 or 1*/*2 in case of monkeys one or two, respectively.

### Data analyses: Neural Activity

#### Firing Rates

The trial-averaged firing rates were estimated by conditioning on *f*_1_. The analysis window, which encompassed the presentation of the first stimulus and the delay period, slid in uniform steps. The analysis intervals were initially positioned at the onset and the offset of *f*_1_, respectively. The temporal window size was set to 250 ms, with a sliding interval of 25 ms, except during the analysis of the terminal states (explained below), where a window of 100 ms with a sliding interval of 10 ms was employed.

#### Mutual Information (MI)

The information that neurons carried about the stimuli was measured using MI [61]. This analysis let us to extract information from the single neuron response, quantified as the number of spikes, *r*_*t*_, fired in the t-th time bin. The quantities *r* = {*r*_1_, …, *r*_*L*_}, where *L* is the total number of bins, were computed using a temporal window of size 250 ms, sliding in steps of 25 ms. *R* collects all values of *r*. If the information sought is about the either the first or second stimulus, all possible values of *f*_*x*_ (where *x* indicates that it could refer to either the first or the second stimulus) are collected in *F*_*x*_. With this, the MI about *f*_*x*_ carried by a neuron is

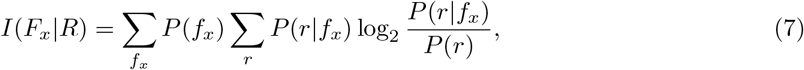

here *P* (*f*_*x*_) is the *f*_*x*_ probability and *P* (*r*|*f*_*x*_) is the conditional probability of observing a response *r* given that the presented stimulus was *f*_*x*_. Equation (7) was computed using a MATLAB-based toolbox given by [62]. The authors also discuss strategies to address the presence of an upward bias in the estimation of MI caused by limited sampling of response probabilities. The strategy we pursued involved an analytical correction combined with a bootstrap procedure. Firstly, the analytic correction, proposed by [63], assumes that the bias correction can be approximated by a second-order expansion in the inverse of the number of trials, *N*_*trial*_,

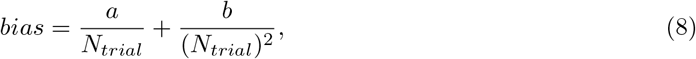

where *a* and *b* are the first two correction terms. However, this correction was not exhaustive as the previous series expansion did not provide a perfect estimate of bias. Hence, we supplemented it with a bootstrap procedure (also included in the toolbox), randomly pairing stimuli and responses to disrupt their correlation. The average value of the resulted bootstrapped MI was used as an estimate of the residual bias. Instead of subtracting this quantity from the corrected information, we used it as a baseline for determining the significance of the corrected MI (SI Appendix, Figs. S1 and S3).

Additionally, the percentage of neurons exhibiting significant MI over time was calculated (Fig. 2; SI Appendix, Fig. S2). In order to compute the significance of this neuron percentage, we employed a shuffling procedure. This involved pooling together the firing rates of all neurons from all trials and carrying out 25 random permutations without replacement. Using this shuffled dataset, we calculated, bin-by-bin, the mean and confidence intervals of the percentage of neurons with significant MI (horizontal shadings in Fig. 2). However, there are numerous isolated instances where the percentage of neurons is marginally above the significance threshold (above the horizontal shading). To assess whether these isolated bins significantly exceed the chance level, we conducted a run-length cluster analysis [56], [57]. This significance evaluation involved determining the proportion of neurons displaying significant MI through 100 random shuffles of the dataset. A null distribution was created by summing the lengths of consecutive significant bins. By using the 95th percentile of this run-length distribution as a significance threshold, we identified which run lengths significantly exceeded the expected chance occurrences (indicated by the lower red and blue data points in Fig. 2; SI Appendix, Fig. S2).

#### Mutual information in error trials

To compute the MI about *f*_1_ in error trials (Fig. 2D), we employed a procedure devised in [39], which relies on the conditional distributions of z-scores from all neurons in the database tuned, either positively or negatively, to *f*_1_ [58]. To combine positive and negative monotonic responses into a unified probability distribution, we assigned labels to the frequencies based on their corresponding z-scores obtained from negatively tuned responses, using the inverse order [39].

In set 1, this entails that the first frequency corresponds to the one eliciting the minimal response (32 Hz), while the last frequency corresponds to the one evoking the maximum response (8 Hz). Following Vergara et al.’s approach, we then defined a “joint frequency” 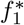 where its minimum value integrates the z-scores associated with *f*_1_ = 8 Hz from positively tuned bins and *f*_1_ = 32 Hz from negatively tuned bins. Finally, we aggregated the normalized activity from all neurons’ trials to construct a probability distribution for each joint frequency, denoted as 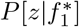 MI was computed using this distribution in an equation analogous to Eq. 7 and corrections for the sampling bias were again taken into account as in Eq. 8. To facilitate comparison, we also assessed the MI in correct trials using the same approach (Fig. 2C).

#### Choice Probability Index

The Choice Probability Index (CPI) evaluates how much the distributions of correct and error responses overlap for each class (*f*_1_, *f*_2_) [59]. The CP index is derived from calculating the area under the receiver-operating characteristic curve (ROC) of the neuron’s firing rate, sorting trials based on the subject’s choice [40]. A value of 0.5 indicates complete overlap, while 1 and 0 indicate entirely separate distributions. A CPI less than 0.5 indicates that the difference between the two distributions is opposite to their expected outcome. Due to the limited number of error trials, we defined the distributions in correct and incorrect trials including the z-scores of all neurons tuned to *f*_1_, similar to our previous description of MI in error trials. These distributions were constructed in each time bin using an equal number of correct and incorrect trials. We assessed CPI at different intervals using a sliding window lasting 150 ms and moving in steps of 25 ms, starting at the onset of *f*_1_ and extending through the presentation of *f*_2_. To gauge the significance of CPI values, we employed a permutation test. This involved randomizing neuronal responses within each time window, essentially reshuffling the correct and error trials. Subsequently, new CPIs were calculated based on this shuffled data. After repeating this process 1000 times, we determined the probability of obtaining CPI values equal to or greater than the initial observations (when CPI exceeded 0.5), or equal to or smaller (when CPI was under 0.5) by random chance using the unshuffled data as a reference [60].

### Data analyses: State-space Analysis and Geometry

#### Principal Component Analysis

We employed principal component (PC) analysis to analyze the firing rates averaged across trials and identify a low-dimensional space that captured a minimum of 90% of the variance in the data. The neural activity, conditioned on the first stimulus, exhibited curved trajectories within this reduced space. The speed associated with these trajectories was computed as the distance covered by a sliding time window of 100 ms, with a 10 ms increment (Fig. 4*C,F*).

#### KiNeT Method for relative distances

We utilized the Kinematic analysis of Neural Trajectory (KiNeT) methodology [13] to determine the relative distance between a pair of trajectories, denoted as *i* and *j*. In this analysis, one trajectory was chosen as the reference (*ref* =*i*), and the corresponding states between the reference trajectory and the other trajectory (*j*) were defined. Specifically, at each time point *t* (aligned to SF1), if the reference trajectory was in state *s*_*ref*_ (*t*), the corresponding state *s*_*j*_(*t*) on trajectory *j* was identified as the one with the minimum Euclidean distance to *s*_*ref*_ (*t*). This allowed us to compute the relative distance *D*_*j*_(*t*) between the two trajectories. For monkey one, the reference trajectory was set to *ref* =22 Hz, while for monkey two, it was *ref* =24 Hz. The relative distances were computed in a PC space capturing at least 90% of the total variance during the presentation of *f*_1_ (Fig. 6*A*; SI Appendix, Fig. S4*A*) and the delay period (Fig. 6*D*; SI Appendix Fig. S4*D*). To estimate the standard errors, we employed bootstrap resampling with replacement, generating 100 resamples. The mean ± 95% confidence intervals (CI) were then calculated from the PC analysis and the corresponding KiNeT analysis performed on each resample.

### Bayesian Model

#### Formulation and Likelihood function

Behavioral data were fitted using a Bayesian model. We assumed that the Bayesian observer has a good understanding of the stimulus set. Specifically, we assumed that: 1) the prior probabilities of the two frequencies correspond to the inherent structure of the set (Fig. 1*C*); 2) the transition probabilities from the first to the second frequency are those derived from the set. In trial *n*, when a frequency *f*_1,*n*_ is presented, we assumed that the Bayesian observer makes an observation (*o*_1,*n*_) of that frequency, defined by a combination of a noisy measurement of *f*_1,*n*_ (denoted as 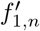)and the observation in the preceding trial (*o*_1,*n*−1_)

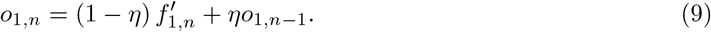

Here *η* is a parameter that measures the contribution of *o*_1,*n*−1_ (if *η* = 0 there is no contribution from sensory history while if *η* = 1 the current stimulus does not contribute). We also assumed that the noisy measurement of *f*_1,*n*_ can be described by a normal distribution

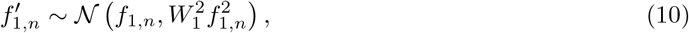

where *W*_1_ is the Weber fraction, a parameter of the Bayesian model to be determined.

In the Results section we observed an absence of transient history effects (pertaining to the previous few trials; see Fig. 2), which leaves us with only the stationary sensory history to consider. From a mathematical standpoint, this implies that in Eq. 9, the observation *o*_1,*n*−1_ is entirely defined by the stationary sensory history of the first stimulus (i.e., it does not contain explicit contributions from recent history), which we assume to be Gaussian. The mean and variance of the stationary variable are obtained by self-consistency arguments (mathematical derivation is provided in the Supporting Text section of the SI Appendix)

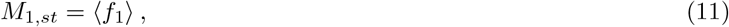

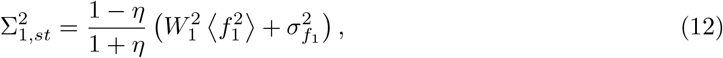

where ⟨*f*_1_⟩, 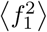 and 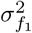 are the mean, the second moment and variance of the probability of the first frequency, 𝒫 (*f*_1_), defined by the experimental set. Within this Gaussian approximation the observation in the current trial is also a Gaussian variable, 𝒪 _1_, with mean ℳ _*n*_ and variance 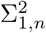,

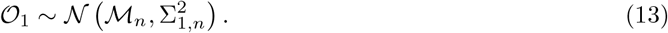

The distribution of this variable defines the likelihood function, that we denote by *P* (𝒪 _1_|*f*_1,*n*_). The mean and variance of the observation probability in the current trial are given by

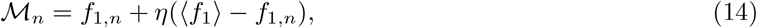

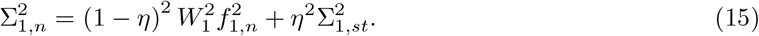

Lastly, we assumed that the second frequency is exclusively influenced by measurement noise. Denoting by *f*_2,*n*_ the value of the second frequency presented in trial *n* and by 𝒪 _2_ the corresponding Gaussian observation, we have

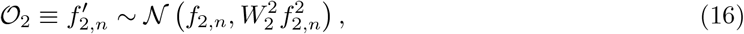

where *W*_2_ is the Weber fraction. In the simulations, the observations 𝒪_1_ and 𝒪_2_ take values ranging from 1Hz to 100Hz, in 1Hz intervals. Note that the observation defined in Eq. 9 serves to incorporate into the model the influence of the stimulus mean, a factor highlighted in previous studies [7], [17], [18], [30], as well as its variance.

#### Beliefs

The computation of posterior probabilities closely aligns with a Bayesian model used for tactile frequency discrimination (Sarno et al., 2022). However, the key distinction lies in the likelihoods, which are determined by the derived expressions mentioned earlier. In our analysis, we represent the discrete prior probabilities of the first frequency as 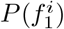, where the index *i* corresponds to the possible values of that frequency within the stimulus set (see Fig. 1*B*). Note that according to our assumptions, the prior probability coincides with the set probability. According to Bayes’ theorem, the posterior distribution 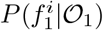 is obtained by combining the noisy observation with the prior distribution

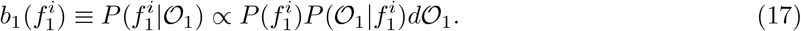

The left hand side indicates that the posterior probability is also known as the belief 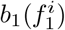 about *f*_1_. Likewise, the posterior probability 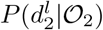, or belief 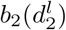 about *f*_2_ combines the noisy observation 𝒪_2_ with its prior

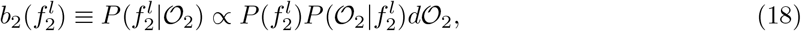

where 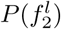 is the discrete prior probability of the second frequency (Fig. 1*C,* *Right*) and *l* labels the possible values of that frequency in the stimulus set (Fig. 1*B*). Note that we should specify whether this prior probability depends on the presented value of the first frequency. We assumed that the noise affecting that frequency is sufficiently large, making this dependence minimal by the time the second stimulus is presented.

Let us denote class *k* as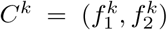, where *k* = 1, …, 10 or 12 (for monkey one or two, respectively) is the class label (Fig. 1*B*). The joint posterior probabilities, or joint beliefs, indicating the likelihood that class *C*^*k*^ was presented in the current trial given the observations𝒪_1_ and 𝒪_2_, can be expressed as follows [10]

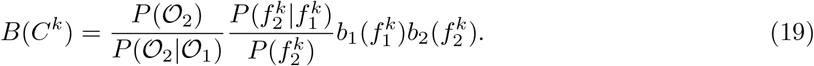

A mathematical derivation of this expression is provided in the Supporting Text section of the SI Appendix. The initial quotient is a normalization factor. 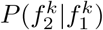 represents the transition probability matrix indicating the probability of 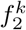 being presented after 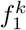 If the animals possess a good understanding of the set’s structure, in the model, the transition matrix elements that are non-zero exclusively correspond to the classes within the set. To verify this assumption, in the Results section we also examined the scenario where transitions not allowed by the stimulus set are given non-zero probabilities (*ϵ*).

The Bayesian observer utilizes the relative beliefs to make a decision between the choices “*f*_1_ smaller” (*S* ≡ “*f*_1_ *< f*_2_”) and “*f*_1_ larger” (*L* ≡ “*f*_1_ *> f*_2_”). These beliefs are calculated by aggregating the class beliefs *B*(*C*^*k*^) for classes *k* associated with a particular choice.

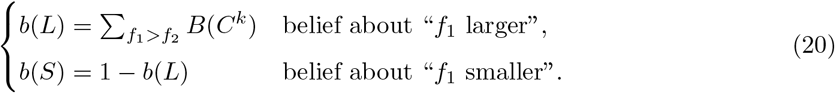

#### Probability of deciding “f_1_ smaller”

Given the information 𝒪_1_ and 𝒪_2_, when the current class is 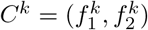, the Bayesian observer selects the option “*f*_1_ smaller” with the probability

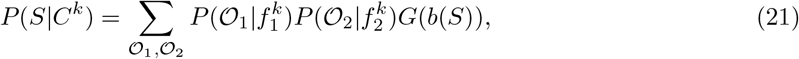

where *G*(*b*(*S*)) equals 1 if *b*(*S*) > 0.5 and and it equals zero otherwise [10], [19].

### Fitting the Bayesian Model to Data

The model was fitted by minimizing mean squared differences between the accuracy curve derived from Eq. (21) and the corresponding behavioral curve obtained from experimental data (Fig. 1*D*). Minimizations were done using MATLAB’s *simulannealbnd* function which implements a simulated annealing algorithm.

### Model comparison

*Akaike Information Criteria* (AIC):

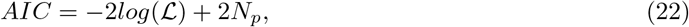

here, the number of model parameters is given by *N*_*p*_. ℒ is the likelihood of obtaining the observed choice (“*f*_1_ *< f*_2_” or “*f*_1_ *> f*_2_”) across all trials:

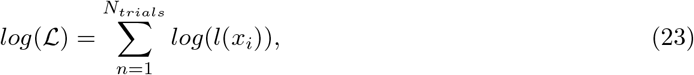

where *l*(*x*_*i*_) is the likelihood function that takes the values *l*(*x* = “*f*_1_ *< f*_2_”) = *P* (“*f*_1_ *< f*_2_”) or *l*(*x* = “*f*_1_ *> f*_2_”) = *P* (“*f*_1_ *> f*_2_”) for a given class and choice.

*Bayesian Information Criteria* (BIC):

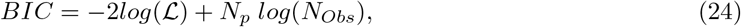

where *N*_*Obs*_ is the number of observations (sample size).

### Estimation of *f*_1_ and its relationship with state-space geometry

To assess whether geometric features of the trajectories encoded an estimate of the first stimulus, we conducted separate comparisons between three estimators: the true *f*_1_, the Bayesian estimator *f*_1,*Bayes*_, and the maximum a posteriori (MAP) estimator *f*_1,*MAP*_. We evaluated these estimators using measures of relative distances during both the presentation of *f*_1_ and the delay period, as well as the terminal states. To accomplish this, we computed the estimators in single trials by simulating the model (using eqs. (14, 15)). Then, we obtained the means *f*_1,*Bayes*_ and *f*_1,*MAP*_ as corresponding averages over trials.

#### Bayesian and MAP estimators

For a given trial *n*, the Bayesian estimator of 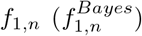 results from a loss function [64]; taking it as the squared error

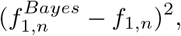

the Bayesian estimator is the mean of the first frequency weighted with the posterior probability at trial 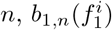

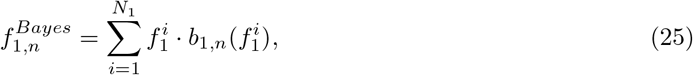

where *N*_1_ represents the number of possible values for *f*_1_ in the set, which is equal to 7 for monkey one and 6 for monkey two.

The MAP estimator 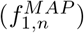 is the value of *f*_1_ that maximizes the belief,

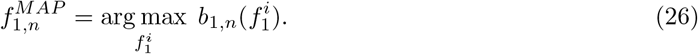

After determining the values of the model parameters, we conducted simulations consisting of 10,000 trials and computed the corresponding Bayesian estimator (Eq. (25)) and MAP estimator (Eq. (26)). Taking the trial averages of these estimators, we obtained *f*_1,*Bayes*_ and *f*_1,*MAP*_, respectively.

#### Mean relative distances

To determine which of the three estimators provided the best explanation for the relative distances between trajectories (see KiNeT section above), we compared the following three hypotheses during the presentation of the first stimulus (Fig. 6*B*; SI Appendix, Fig. S4*B*) and the delay period (Fig. 6*E*; SI Appendix, Fig. S4*E*):

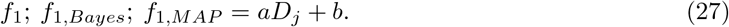

Here *D*_*j*_ (*j* = 1, …, *N*_1_. Again, *N*_1_ = 7 in case of monkey one and *N*_1_ = 6 in case of monkey two) is the temporal mean of the relative distances [14] during either the presentation of the first stimulus, 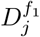, (inset from Fig. 6*A*; SI Appendix, Fig. S4*A*) or the delay period, 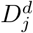, (inset from Fig. 6*D*; SI Appendix, Fig. S4*D*). For both intervals, we utilized an analysis window of 250 ms with a sliding interval of 25 ms. We retained the number of PCs necessary to explain a minimum of 90% of the variance.

#### Distances between states at the end of the delay period (terminal states)

We defined the line of terminal states as the line that results from connecting all consecutive terminal states. To examine which estimator is encoded by them, we evaluated the Euclidean distance along the line of terminal states, 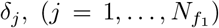 between the final state of each *f*_1_ trajectory and the final state of the trajectory with the smallest *f*_1_ (Fig. 6*G*; SI Appendix, Fig. S4*G*). Note that these distances are independent of the reference state. We then made hypotheses analogous to those in Eq. (27). This test (Fig. 6*H*; SI Appendix, Fig. S4*H*) took place in the last 100 ms before the end of the delay period. The terminal states were considered to be embedded in a 2-dimensional subspace, which accounted for 84% of variance in monkey one and 93% of variance in monkey two.

In the three tests the goodness of the fits was compared using their Root Mean Square Errors (RMSEs; Figs. 6*C,F,I* for monkey one; SI Appendix, Figs. S4*C,F,I* for monkey two). Significance was assessed using a bootstrap procedure. We generated 100 resamples by randomly sampling with replacement from the original neural population. Subsequently, we conducted the tests on each resample and calculated the respective dispersions.

## Code and Data Availability

Matlab codes, as well as the raw data necessary for full replication of the figures will be available upon publication.

## Acknowledgements

This work was partly supported by the grant PGC2018-101992-B-I00 from the Spanish Ministry of Science, Innovation and Universities. We thank Athena Akrami, Rubén Moreno and Alfonso Renart for their valuable insights, which have significantly enhanced the quality of our manuscript.

## Author contributions

N.P. and R.R. designed research. L.S.-F. implemented the numerical computations and analyzed the data. M.B. and N.P. developed the Bayesian model. L.S.-F. and N.P. wrote the paper. M.B. and R.R. revised the text.

## Supporting Information

### Supporting Information Text

#### Methods

##### Bayesian Model. Mathematical derivation of the stationary regime by self-consistency arguments

The objective of this Supporting Text is to obtain the stationary regime (Equations 11 and 12 in the Methods section) using self-consistency arguments. Equation 9 in the Methods section describes the observation of the first frequency in a trial *n* (*o*_1,*n*_) as a linear combination of the current-trial noisy measurement of the first frequency 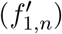 and the observation in the preceding trial (*o*_1,*n*−1_). Therefore, it involves three different stochastic variables, related through a linear expression of the form

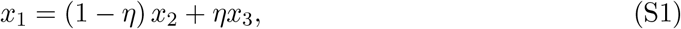

where *x*_*i*_ for *i* = 1, 2, 3 represent random variables, with mean and variance 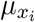 and 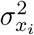 The mean and variances of the random variables are therefore inter-related

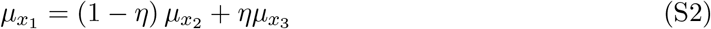

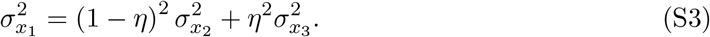

By construction, two of the random variables in Equation 9 of the Methods have the same statistics: the observations in trial *n* and trial *n* − 1. We refer to the mean and variance of the observations using *M*_1,*st*_ and 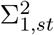. The probability distribution of the remaining random variable, 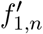, is given by a mixture of Gaussians that depends on the stimulus set. The probability density function of the mixture of Gaussians reads

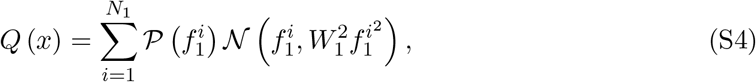

where 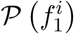 indicates the probability of the first frequency as defined by the stimulus set (Fig. 1*B*), and *N*_1_ represents the number of possible values that first frequency can take within the set (see Fig. 1*B*), which is equal to 7 for monkey one and 6 for monkey two. Therefore, the mean and variance of 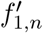, determined by *Q*, can be written down explicitly as

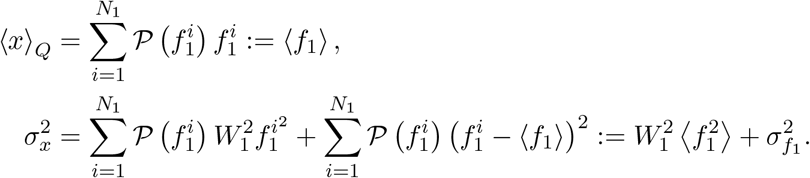

Finally, mapping the identities in Equations S2 and S3 to Equation 9 in the Methods, we obtain

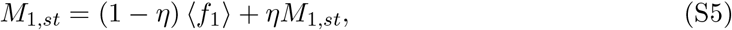

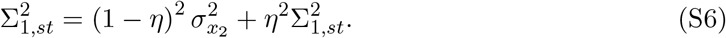

Rearranging terms in the equations above, we lead to Equations 11 and 12 in the Methods section.

##### Bayesian Model. Mathematical derivation of the joint belief (belief on the class)

The aim of this Supporting Text is to derive the expression for the joint belief (Equation 19 in the Methods section) following the mathematical approach outlined in [1]. Denoting class *k* as 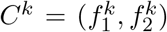, where *k* = 1, …, 10 or 12 (for monkey one or two, respectively) is the class label (Fig. 1*B*), the joint posterior probabilities, or joint beliefs, can be expressed as the likelihood that class *C*^*k*^ was presented in the current trial given the observations 𝒪 _1_ and 𝒪 _2_

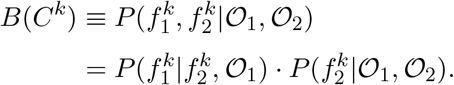

Applying Bayes’ theorem in both factors, the joint beliefs read

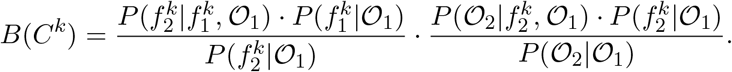

Then, considering that 𝒪 _1_ only depends on the first frequency, and applying again Bayes’ theorem on 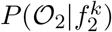, we obtain

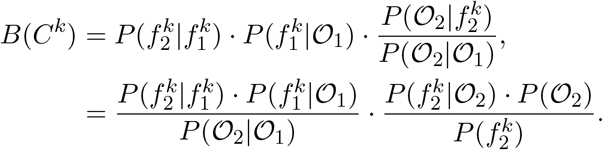

Finally, in terms of the individual beliefs 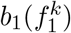 and 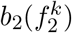 the joint belief can be expressed as follows (Eq. 19 in the Methods section)

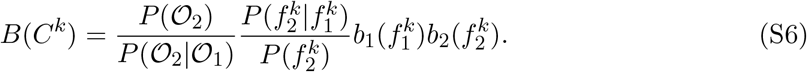

### Supplementary Figures

**Fig. S1.**
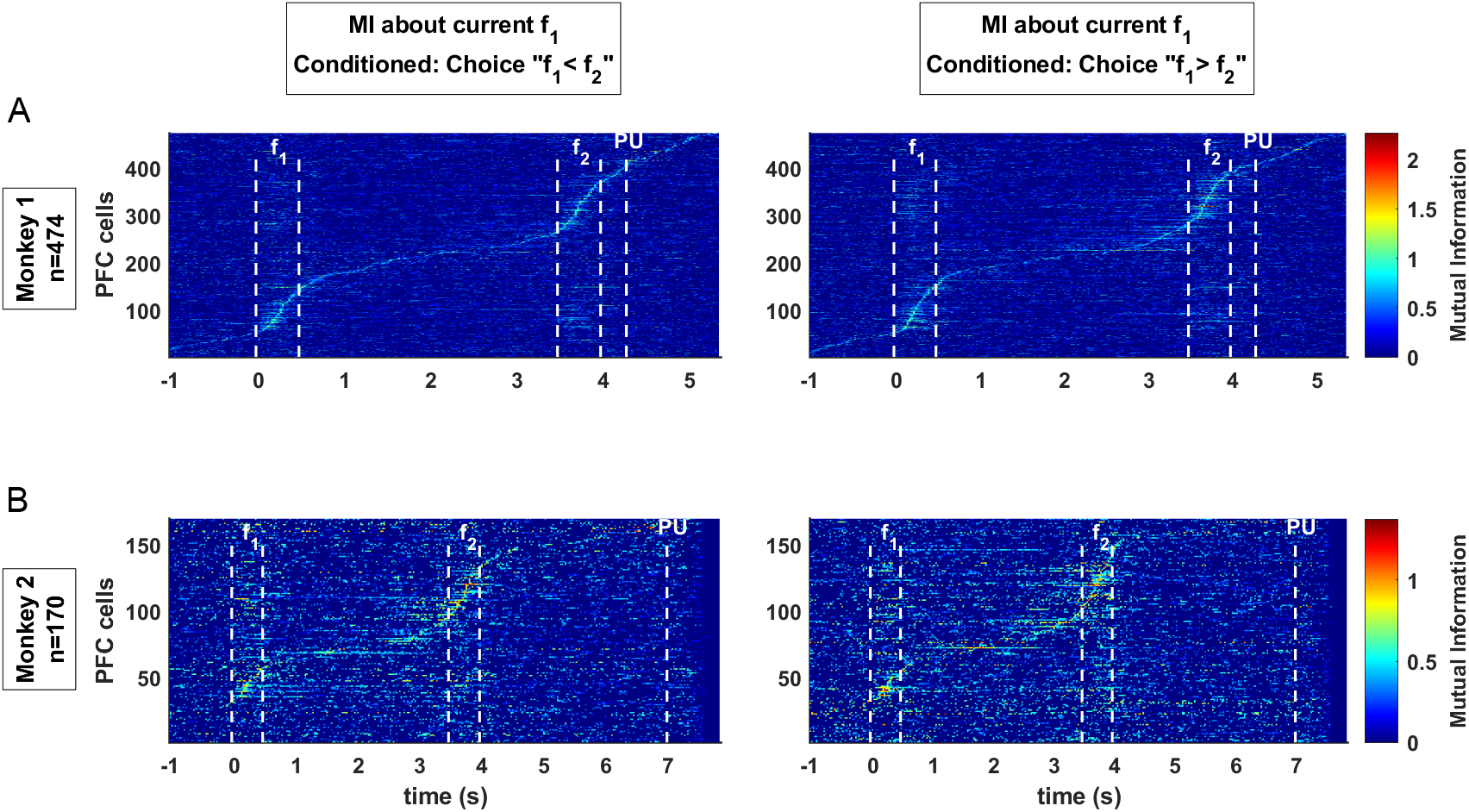
Mutual information (MI) about *f*_1_ in the current trial. MI as a function of time from either *(A)* the 474 cells of monkey one or *(B)* the 170 cells of monkey two and conditioning on rewarded current trials either with choice “*f*_1_ *< f*_2_” *(Left)* or with choice “*f*_1_ *> f*_2_” (*Right)*. Note that this effectively fixes the class in the current trial. Dark blue indicates the non-significant MI values (see Methods).

**Fig. S2.**
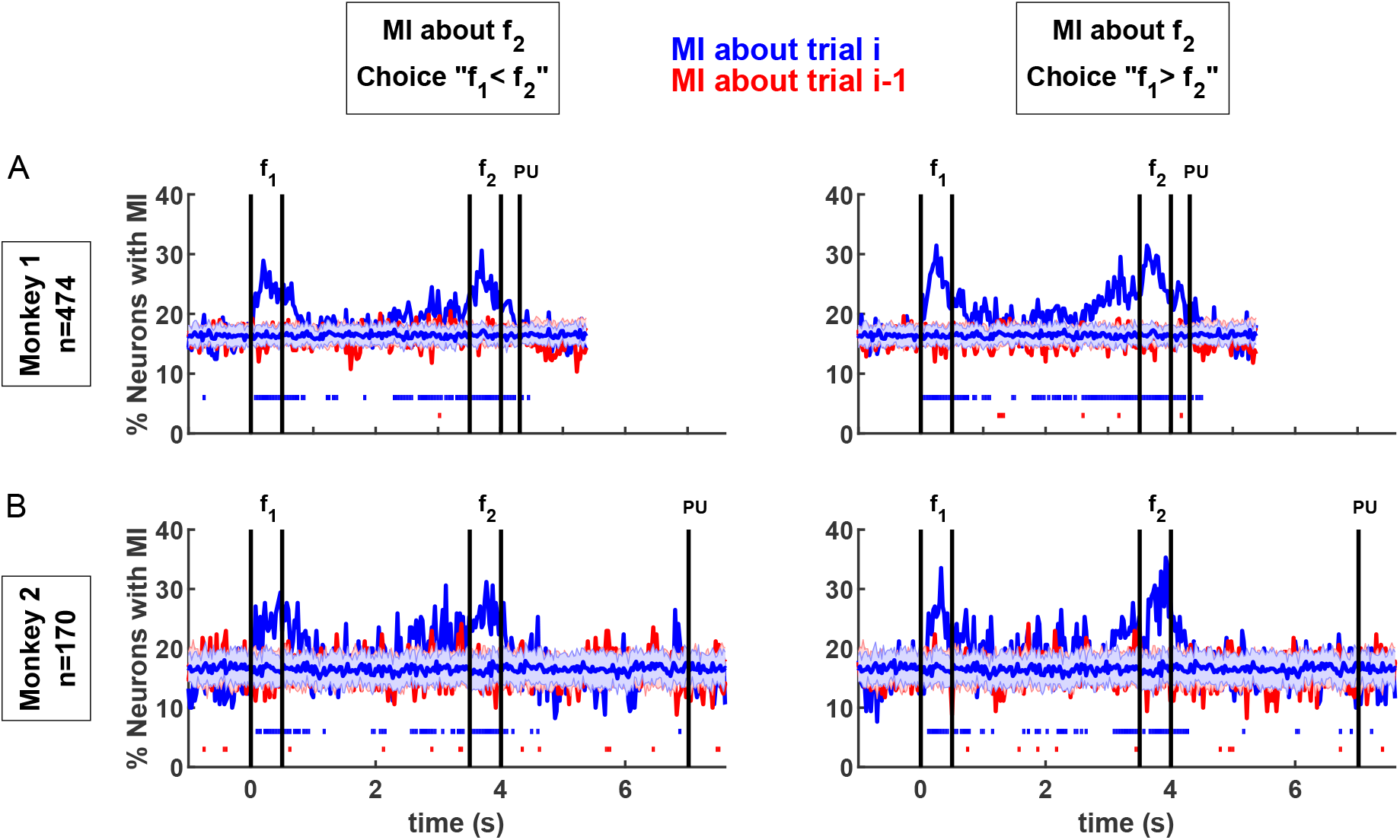
PFC neurons exhibited significant mutual information (MI) about the current trial *f*_2_ but not on the previous trial *f*_2_. *(A)* Percentage of neurons from monkey one with significant MI about the current (blue line) and previous (red line) second stimulus, conditioning on correct and fixed choice (*Left*: choice “*f*_1_ *< f*_2_”; *Right*: choice “*f*_1_ *> f*_2_”) current trials. Note that this effectively fixes the class in the current trial. Horizontal blue and red shadings refer to the mean and standard deviation computed from shuffled data (see Methods) for current and previous trials, respectively. The horizontal lines of dots at the bottom indicate runs of consecutive time bins where a significant percentage of neurons exhibit significant MI, assessed through run length analysis (see Methods). *(B)* Same MI analysis for monkey two.

**Fig. S3.**
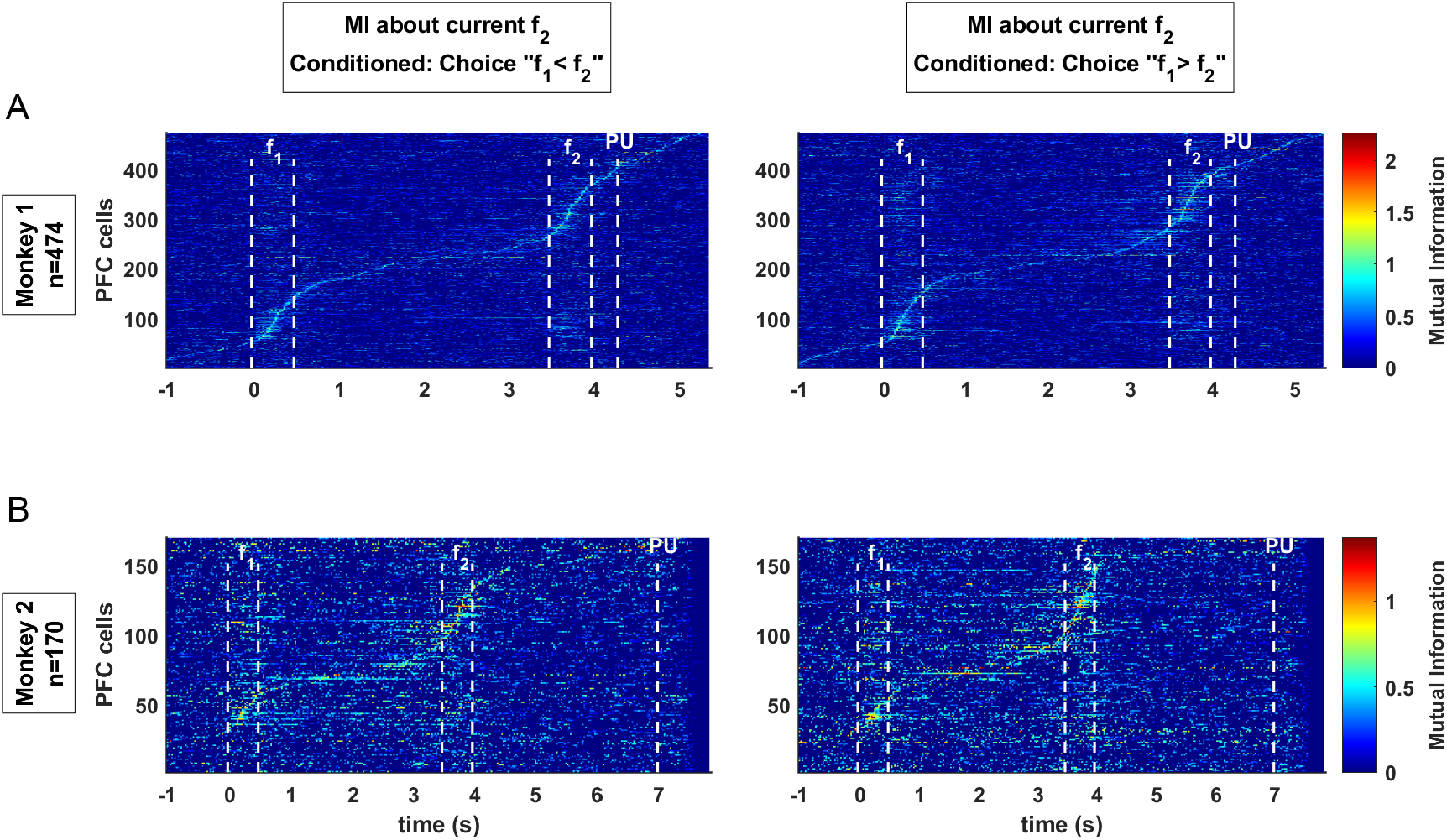
Mutual information (MI) about *f*_2_ in the current trial. MI as a function of time from either *(A)* the 474 cells of monkey one or *(B)* the 170 cells of monkey two, conditioning on rewarded current trials either with choice “*f*_1_ *< f*_2_” *(Left)* or with choice “*f*_1_ *> f*_2_” (*Right)*. Note that this effectively fixes the class in the current trial. Dark blue indicates non-significant MI values (see Methods).

**Fig. S4.**
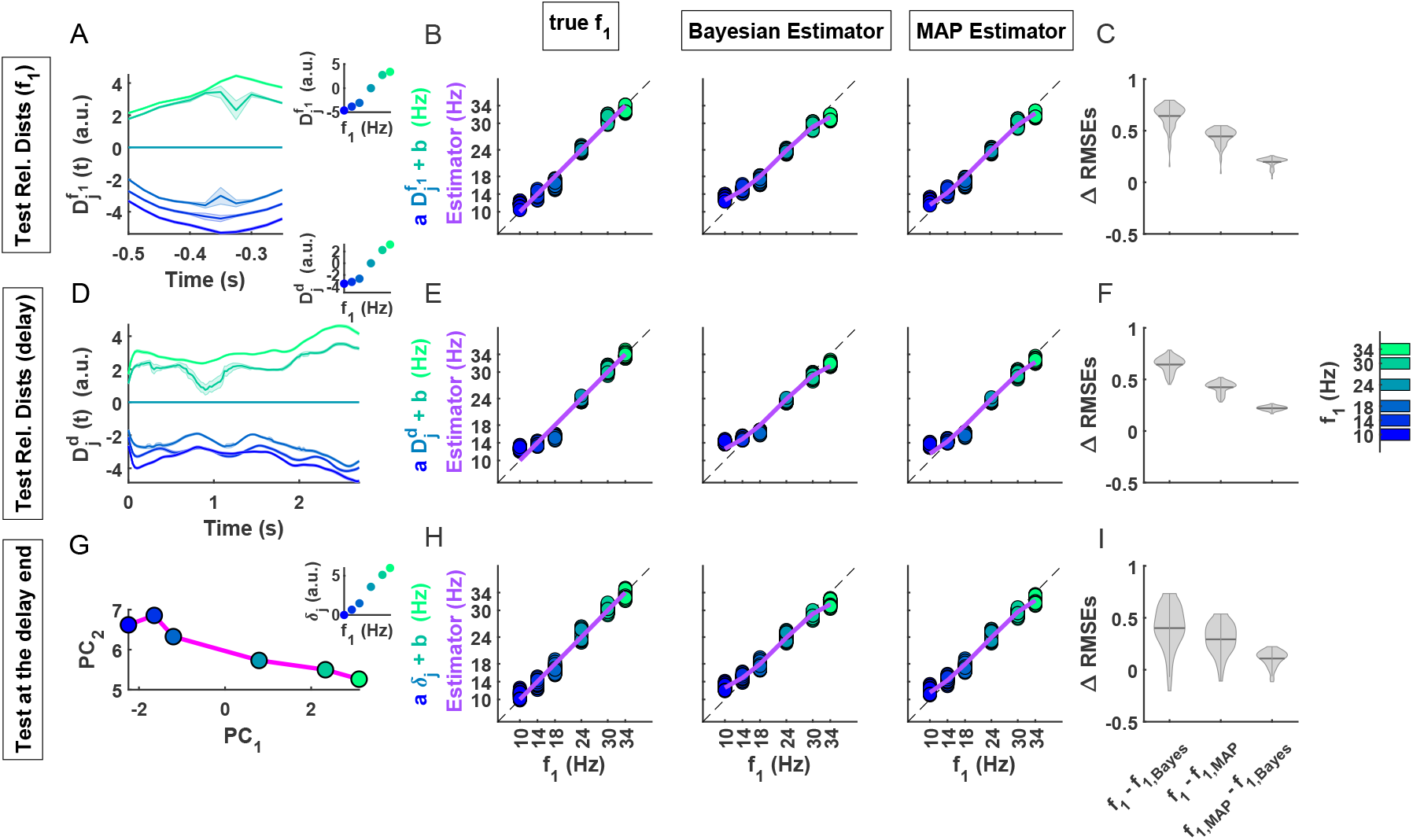
Several geometric features from monkey two code the Bayesian estimator. *(A-C)* KiNeT trajectories and activity-behavior relationship during the presentation of the first frequency. *(A)* Relative distance 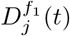, between each trajectory *j* = 1, …, 6 and the reference trajectory *f*_1_=24Hz using KiNeT analysis. Alignment is located at the offset of the first stimulus. The state space that embeds these neural trajectories is constructed using 5 PCs, which account for 91% of the variance. Shadings were built from the mean ± 95% CIs of 100 bootstrap resamples. *Inset* displays 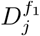, the temporal means of relative distances as function of *f*_1_. *(B)* Linear regression of the mean relative distances 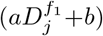 to the true *f*_1_-value *(Left)*, the Bayesian estimator *(Middle)* and the MAP estimator *(Right)* as a function of *f*_1_. Colored dots represent multiple estimates of the regression obtained through bootstrapping (100 resamples). Purple solid lines indicate the true *f*_1_ *(Left)*, its Bayesian estimator *(Middle)* and its MAP estimator *(Right)*. The unity line is represented by a black dotted line. *(C)* Violin plots display the empirical probability density of ΔRMSEs on the y-axis, which represents the differences in RMSEs between the three hypotheses (*f*_1_, Bayesian estimator, and MAP estimator) indicated on the x-axis. A positive ΔRMSE value indicates that the second of the two estimators fits the geometric feature better. The Bayesian estimator fits the data better than the alternative hypotheses for all tests. The width of the gray area represents the density of the shufflings in that range. The horizontal line within each violin plot indicates the mean of the distribution. *(D-F)* KiNeT trajectories and activity-behavior relationship during the delay period. Same format as *A-C*, as well as same alignment at the offset of *f*_1_. Activity analysis with KiNeT was, as well as same alignment at the offset of *f*_1_build from a state space of 14 PCs explaining the 90% of variance. Again, Bayesian hypothesis is chosen by the 100 resamples. *(G-I)* Terminal states and activity-behavior relationship at the end of the delay period. *(G)* Line of terminal states (magenta line and circles) embedded within a 2-dimensional subspace (93% of variance). *Inset* shows the Euclidean distances along the magenta line, *δ*_*j*_, between each *f*_1_ terminal state and the smallest *f*_1_ (taken as reference), as function of *f*_1_. *(H-I)*. activity-behavior relationship in the same format as *B-C*. Here the 90% of resamples choose Bayesian hypothesis.

### Supplementary Tables

**Table S1.** The model parameter *η* specifically regulates the long-term sensory history. The fitted behavioral data correspond to monkey one, featuring 3-second delay period in the first column and 6-second delay period in the subsequent columns. The first and second columns display the fitting of measurement uncertainties (*W*_1_ and *W*_2_) along with *η*. The third column depicts the fitting while only adjusting the *η* parameter (with *W*_1_ and *W*_2_ fixed using values obtained from the 3-second delay data fitting). In the fourth column, adjustments are made to both *W*_1_ and *η* (with *W*_2_ fixed using the value obtained from the 3-second delay data fitting). The RMSE is the root mean square error between the behavioral and model performances. It is noted that the *η* parameter in the last three columns experiences minimal changes. The AIC and BIC criteria (see Methods) applied to the 6-second task fits tend to favor the model where all three parameters are freely adjusted.

|  | **3s Delay** | **6s Delay** | **6s Delay**<br>**(fixed $W_1$ and $W_2$)** | **6s Delay**<br>**(fixed $W_2$)** |
| --- | --- | --- | --- | --- |
| $W_1$ | 0.064 | 0.057 | 0.064 | 0.085 |
| $W_2$ | 0.088 | 0.110 | 0.088 | 0.088 |
| $\eta$ | 0.509 | 0.634 | 0.639 | 0.630 |
| *RMSE* | 0.020 | 0.028 | 0.031 | 0.029 |
| *BIC* | 4547 | 1267 | 1282 | 1287 |
| *AIC* | 4525 | 1250 | 1276 | 1275 |

**Table S2.**
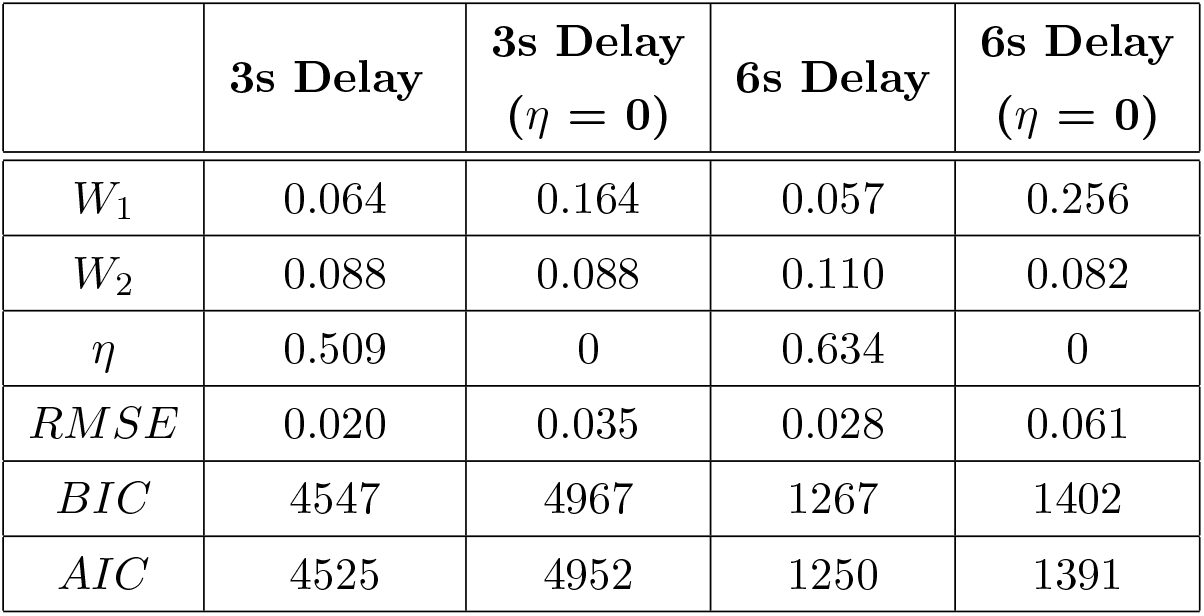
Comparison between the Bayesian model with non-zero *η* and the model with *η* = 0. Fitted behavioral data for monkey one with both 3-second and 6-second delay periods. *W*_1_ and *W*_2_ are the Weber fractions of the first and second stimulus, respectively. The parameter *η* regulates the long-term sensory history. RMSE denotes the root mean square error, measuring the difference between the behavioral data performance and the model performance. RMSE values and the criteria AIC and BIC (see Methods) tend to favor the model incorporating knowledge about the long-term sensory history.

**Table S3.** Comparison between the Bayesian model with non-zero *ϵ* and the model with *ϵ* = 0. Fitted behavioral data for monkey one (M1) with both 3-second and 6-second delay periods, and for monkey two (M2). *W*_1_ and *W*_2_ are the Weber fractions of the first and second stimulus, respectively. The parameter *η* regulates the long-term sensory history. The parameter *ϵ* introduces non-zero probabilities for transitions not allowed by the stimulus set. RMSE denotes the root mean square error, measuring the difference between the behavioral data performance and the model performance. As shown in the table, the model extended with the *ϵ* parameter does not result in any substantial changes in the other parameters compared with the simpler model, with the *ϵ* value remaining very close to zero.

|  | **M1:**<br>**3s Delay** | **M1: 3s Delay**<br>**(fitting $\epsilon$)** | **M1:**<br>**6s Delay** | **M1: 6s Delay**<br>**(fitting $\epsilon$)** | **M2** | **M2**<br>**(fitting $\epsilon$)** |
| --- | --- | --- | --- | --- | --- | --- |
| $W_1$ | 0.064 | 0.052 | 0.057 | 0.056 | 0.054 | 0.062 |
| $W_2$ | 0.088 | 0.081 | 0.110 | 0.110 | 0.108 | 0.108 |
| $\eta$ | 0.509 | 0.516 | 0.634 | 0.632 | 0.430 | 0.420 |
| $\epsilon$ | 0 | 0.0025 | 0 | 0.0003 | 0 | 0.0283 |
| $RMSE$ | 0.020 | 0.020 | 0.028 | 0.028 | 0.019 | 0.019 |

